# Histone diversity in the archaeal domain of life

**DOI:** 10.1101/2025.06.26.661778

**Authors:** Shawn P. Laursen, Karolin Luger

## Abstract

Archaea represent a distinct domain of life that is genetically and biochemically unique from bacteria and eukaryotes. Two-thirds of all archaea encode histones, proteins that are ubiquitously used to structure chromatin in eukaryotes. Archaeal histone sequences are much less conserved than their eukaryotic counterparts, yet insight into how they structure DNA is limited to only a few species that fail to represent the diversity of the archaeal domain. Archaea have adapted to the most diverse and extreme environments on our planet, requiring protection of the genome against a multitude of external pressures. Here, we use bioinformatics, structure prediction, and molecular dynamics simulations to survey the diversity of histone-like sequences in all available archaeal genomes and to understand how they might interact with DNA. We have identified five distinct types of histones which are combined in seven different strategies, involving either single histones, multiple histones of the same type, or combinations of several types of histones in one genome. We show that some strategies correlate with environmental pressures, and some are phylogenetically restricted. Despite highly divergent amino acid sequences, structure predictions and simulations suggest similar histone DNA binding modes for most classes. Our work provides a guide to efficiently survey diverse strategies for histone-based DNA organization in archaea using biophysical and structural approaches, for a complete view of the rich diversity of histone strategies in the archaeal domain in a targeted manner.

## Introduction

Histones are small, highly basic proteins consisting of three α helices connected by two short loops (the ‘histone fold’) that form either homo- or heterodimers via a ‘handshake motif’ ^1,2^. These proteins are present in genomes across all domains of life and are also found in some viruses^3–8^. The best-studied histones are from the eukaryotic domain where heterodimers of histone H2B-H2A and H3-H4 assemble into an octamer that wraps 147 base pairs of DNA to form nucleosomes^3^. Eukaryotic histones are highly conserved and ubiquitously present across the entire domain. Homologues of the four types of histones also are encoded in the genomes of several ancient double-stranded DNA viruses that infect amoeba^9,10^. The amino acid sequences of these histones are rather divergent amongst giant viruses, and differ in many ways from those of eukaryotes. While the overall topology of nucleosomes reconstituted from these viruses appears to be conserved (at least for the two distantly related viruses where this has been studied^6, 7,11^), the individual histone chains can be found in a variety of tandem, triple, and even quadruple combinations and in truncated forms in different viruses^10^.

A subset of bacteria also have histone-like proteins, which were likely acquired through horizontal gene transfer^12^. While some of these are attached to other domains of mostly unknown function, many are standalone histones that are abundantly expressed and associated with the nucleoid^5^. Only two of these putative bacterial histones have been studied in detail and their interaction with DNA is markedly different to that of eukaryotic histones. Histones from *Bdellovibrio bacteriovorus* create long protein-coated DNA filaments through ‘edge-on’ binding rather than wrapping the DNA to form discrete nucleosomes, although the binding mode on longer DNA is somewhat controversial^5,13,14^. A recent preprint suggests yet another binding mode for a histone encoded by *Leptospira perolatii*^15^. Clearly, more research is needed to understand how histones are used in bacterial genome organization.

Histones are widespread in the domain of archaea. They were first discovered by John Reeve and coworkers in 1990^16^, and we now know that the majority of archaeal genomes encode at least one type of histone. As more archaeal species are discovered at a rapid rate through advances in genomic sequencing, there is evermore diversity to consider^17–19^. Archaeal histones exhibit much more sequence divergence than their eukaryotic counterparts^20^, which are amongst the most conserved proteins known^21^. Because archaea are found in many different and often punishing environments, their proteins must have evolved to cope with extreme conditions^22–26^. Unlike in bacteria, where histone genes are sparse, histones seem to be a deeply rooted feature of archaea, occurring in most higher taxa, with the notable exception of *Thermoplasmata* (formerly *Crenarchaeota*)^26–28^. A select few archaea encode histones with tails, with the potential for post-translational modifications. These organisms are mainly from the Asgard phylum, which are thought to be most closely related to eukaryotes^20,29^.

At least two closely related hyperthermophilic archaea, *Thermococcus kodakarensis* and *Methanothermus fervidus*, have histones that package DNA into so-called ‘hypernucleosomes’, slinky-like assemblies where the geometry of the DNA superhelix closely mimics the superhelix formed by stacked eukaryotic nucleosomes, using near-identical features of the histones to engage the DNA backbone^4,30^. To date, research into archaeal histone-DNA complexes is limited to these two organisms (but see a recent preprint article^31^). In *T. kodakarensis*, histones contribute to transcription regulation^4^. Additional studies utilizing molecular modeling of sequences from methanogenic archaea, and ChIP-seq in *Halobacterium salinarum* have begun to shed light on the function of histones in these organisms^32,33^. Here, we parsed the diversity of histone sequences in archaea by mining predicted proteins databases. We grouped archaeal histones into five major clusters based on four biophysical properties (length, isoelectric point, hydrophobicity, and instability index). We then identified seven strategies by which different organisms combine histones; employing either a single histone, or various combinations of histones in one genome. To understand possible co-dependencies between histones, we analyzed the seven strategies separately, to allow us to tease apart, for example, whether basic histones that occur as the only histone in an organism have different features compared to those that co-exist with other basic or acidic histones. We predicted the structure of the main histone combinations and inferred their ability to bind DNA using molecular dynamic simulations, providing a starting point for targeted structural and biophysical analysis.

## Results

### Archaeal histones can be grouped into five clusters

We first identified putative histones in the predicted proteomes of all 5,869 available archaeal genomes in release 220 of the Genome Taxonomy Database (GTDB)^34^. Protein coding sequences in this database were predicted from single genomic assemblies representing unique species. Metadata including sampling location, genome size, and GC content were also calculated or collected. To identify histone sequences, we used HMMSearch with archaeal (PF00808) and eukaryotic (PF00125) histone PFAM models. We tested a variety of search strategies using different HMMer tools with a range of stringency cutoffs and found that HMMSearch with a liberal stringency captured most of the diversity found in the sequence space, without adding too much noise (Supplemental Figure 1).

We then applied DBSCAN (Density-Based Spatial Clustering of Applications with Noise) to perform unsupervised clustering of the presumptive histone sequences, using the four easily calculated physical parameters with the most variance: length, instability index, isoelectric point (pI), and hydrophobicity/GRAVY score. Full clustering details can be found in the Methods. Briefly, we optimized the clustering parameters using small randomly sampled datasets, extracted the physical parameters that define each cluster, and used those bounds to label proteins in the overall dataset (Supplemental Table 1).

We used these physical parameters to group histones into five distinct clusters of histone-like proteins (Figure 1a): basic singlets (cluster 1, blue), acidic singlets (cluster 2, red), acidic doublets (cluster 3, orange), acidic ‘miniatures’ (cluster 4, yellow), and acidic quadruplets (cluster 5, green). We selected the centroid sequence from each cluster and predicted their structures with AlphaFold3 (Figure 1b, Table 1). Basic and acidic singlets form the characteristic histone fold that resembles the experimentally determined structure of the basic singlet HMfB (pdb 1A7W). Cluster 3 histones comprise two histone fold domains that are linked in a single polypeptide chain (colored in black in Figure 1b), and they are predicted to form a structure that resembles the HMfB homodimer (pdb 1A7W). The acidic miniature histone (cluster 4) is predicted to have an α2 helix that is shortened by one turn and also has a very rudimentary α3 helix, and in this it resembles the bacterial histone Bd0055 (pdb 8VVX), although the latter is positively charged overall. Finally, cluster 5 histones are unusual in that they consist of a long acidic chain with four predicted histone fold motifs. Archaeal histones have a bimodal distribution in terms of their charge: overall, a surprisingly large percentage (26.7 %) of the 7,157 predicted histones are acidic in character, while histones with neutral charge are largely absent (Figure 1c).

**Figure 1:**
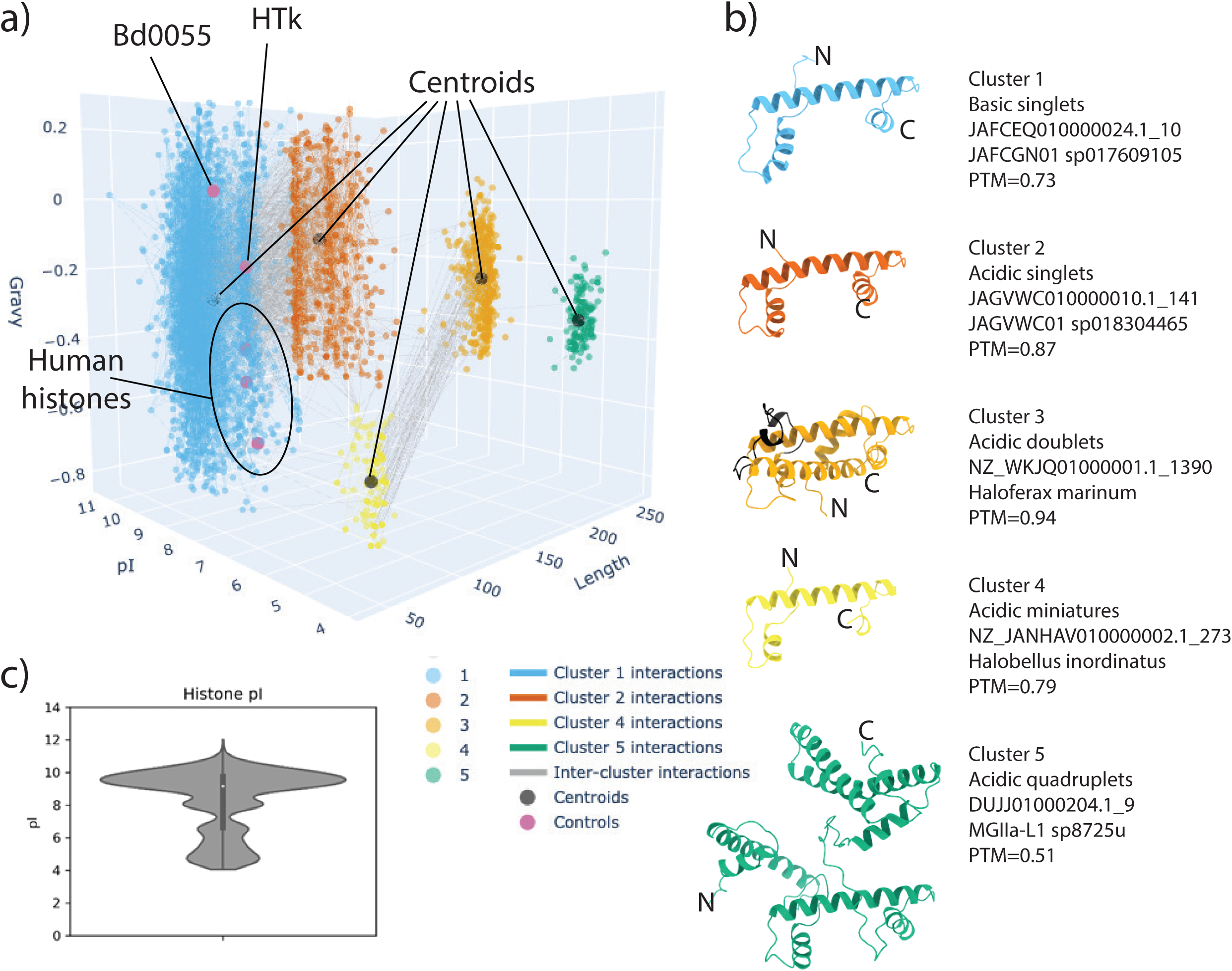
Archaeal histones can be grouped into five clusters. **a)** Clustering of 6,473 archaeal histone sequences using DBSCAN, plotted by length (number of residues), pI (isoelectric point), and GRAVY score (hydrophobicity). Sequences were clustered using these three dimensions plus a fourth, the instability index (not shown). Sequences closest to the center of each cluster in the four dimensions are denoted with a black dot; gene names for the centroids are listed in b). Control histones (human histones H2A, H2B, H3, and H4, bacterial histone Bd0055, and archaeal histone HTkA) are indicated by purple dots. An interactive version of this figure can be found in supplemental materials, all sequences are listed in supplemental spreadsheets. **b**) AlphaFold3 structure predictions of single chains of the centroids determined in **a),** along with defining characteristic, gene name, genome, and AlphaFold3 PTM (confidence) score. N and C-termini are indicated. The linker region connecting the two histone fold domains in cluster 3 is shown in black. **c)** Distribution of isoelectric points of all archaeal histones shown in a). A multimodal distribution can be observed around pI values of 10, 8, 6.5 and 4.5. 26.7 % have an acidic isoelectric point. Isoelectric points around neutral are under-represented.

**Table 1.**
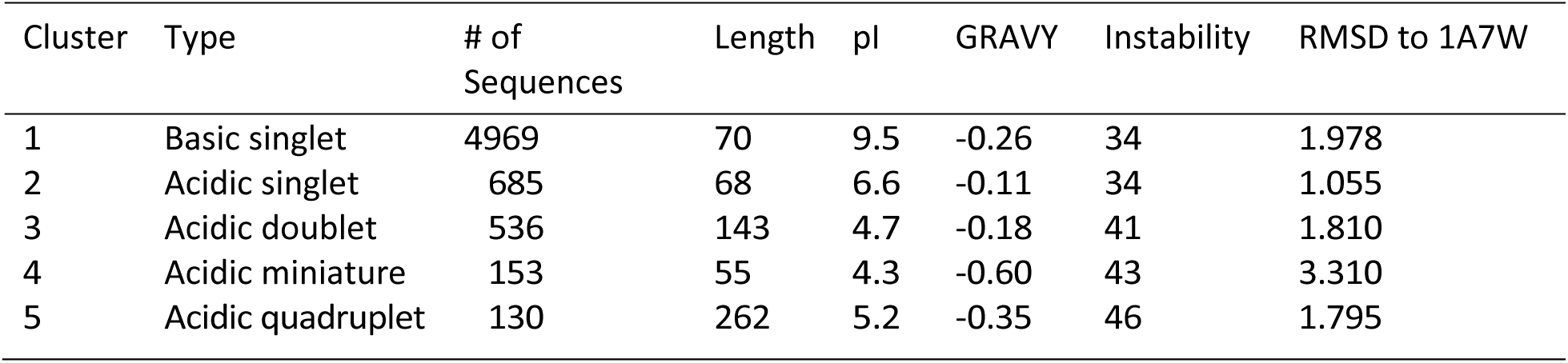
Median values of physical parameters of histone clusters. Number of sequences in each cluster, and median values for length of protein (amino acids), isoelectric point (pI), hydrophobicity (GRAVY score), instability index, and RMSD to histone HMfB from pdb 1A7W to the best aligning histone fold (Å) for each histone cluster were calculated after clustering in Figure 1. See Supplemental Table 2 for ranges of values of each cluster.

| Cluster | Type | # of Sequences | Length | pI | GRAVY | Instability | RMSD to 1A7W |
| --- | --- | --- | --- | --- | --- | --- | --- |
| 1 | Basic singlet | 4969 | 70 | 9.5 | -0.26 | 34 | 1.978 |
| 2 | Acidic singlet | 685 | 68 | 6.6 | -0.11 | 34 | 1.055 |
| 3 | Acidic doublet | 536 | 143 | 4.7 | -0.18 | 41 | 1.810 |
| 4 | Acidic miniature | 153 | 55 | 4.3 | -0.60 | 43 | 3.310 |
| 5 | Acidic quadruplet | 130 | 262 | 5.2 | -0.35 | 46 | 1.795 |

### Histones are used in seven different strategies in the archaeal domain

According to our cutoff, of the 5,869 archaeal genomes in the GTDB, 3,931 (67%) encode at least one putative histone (Figure 2a). Because each sequence represented in Figure 1a is associated with a unique species, we were able to determine which genomes encode more than one histone and which combinations are the most prevalent. About 60% of all histone-encoding genomes have only one single histone gene from either cluster 1, 2, 3, or 5 (Figure 2a, b) and species encoding more than three histones are rare. We classify the genomes encoding only one single histone from a specific cluster as ‘single 1,2,3 or 5’, to separate them from genomes which contain different combinations of histone that also may include histones from the same clusters. Among genomes harboring more than one histone, genomes containing two or more histones from cluster 1 (basic singlets) are the most prevalent strategy, termed ‘multiple 1’ (the model organism *M. fervidus* is an example for this strategy). We also observe combinations of representatives from clusters 1&2 and clusters 3&4, termed combination 1&2 or combination 3&4, respectively (Figure 2b, Supplemental Figure 2). Representatives from cluster 4 (acidic miniatures) are almost always paired with an acidic singlet (cluster 3), and cluster 5 histones always occur as the sole histone-encoding gene. To simplify our analysis, we focused on general trends and restricted our further analysis to these seven most prevalent combinations of histones (single 1,2,3 and 5; multiple 1; combination 1&2; and combination 3&4) which represent *>*98% of histone-encoding archaeal genomes (indicated by a line in Figure 2b).

**Figure 2:**
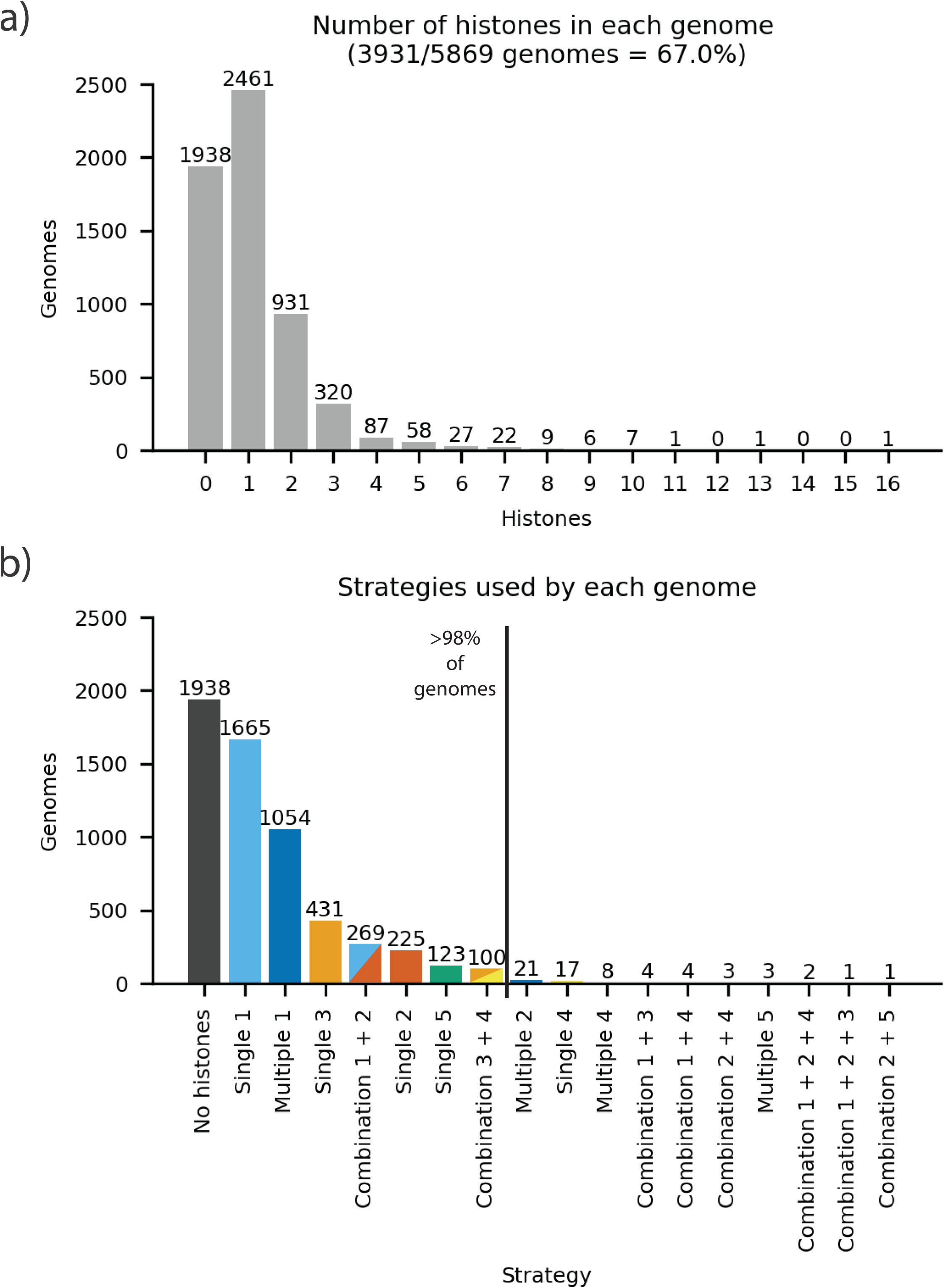
Histones are employed in seven major ‘histone strategies’ in archaea. **a)** Distribution of predicted histone genes per genome. The majority of genomes contain three or fewer histones, and 33 % of genomes encode no clearly identifiable histone (using our relatively conservative cutoff). **b)** Distribution of genomes utilizing a particular histone strategy. These include genomes with only a single histone, genomes with multiples of the same type, or those with combinations of more than one type. Histone clusters are colored as in Figure 1. This study focuses on strategies that are represented by 100 or more genomes (indicated by vertical line)

### Some strategies are taxonomically restricted

Histones that fall into cluster 1 (basic singlets) are widely dispersed across the entire domain of archaea and indeed seem to represent the typical archaeal histone (Figure 3a). They occur either as the sole histone or in combination with other basic singlets throughout the domain. Acidic singlets (cluster 2) are also pervasive, either as the only histone in the genome, or paired with a basic singlet. In contrast, histones from clusters 3, 4, and 5 are phylogenetically restricted to specific taxa. In particular, histones from cluster 3 (acidic doublets) are mostly restricted to the class of Halobacteria, while representatives of cluster 5 (acidic quadruplets) are exclusive to members of the order Poseidonales (Figure 3b).

**Figure 3:**
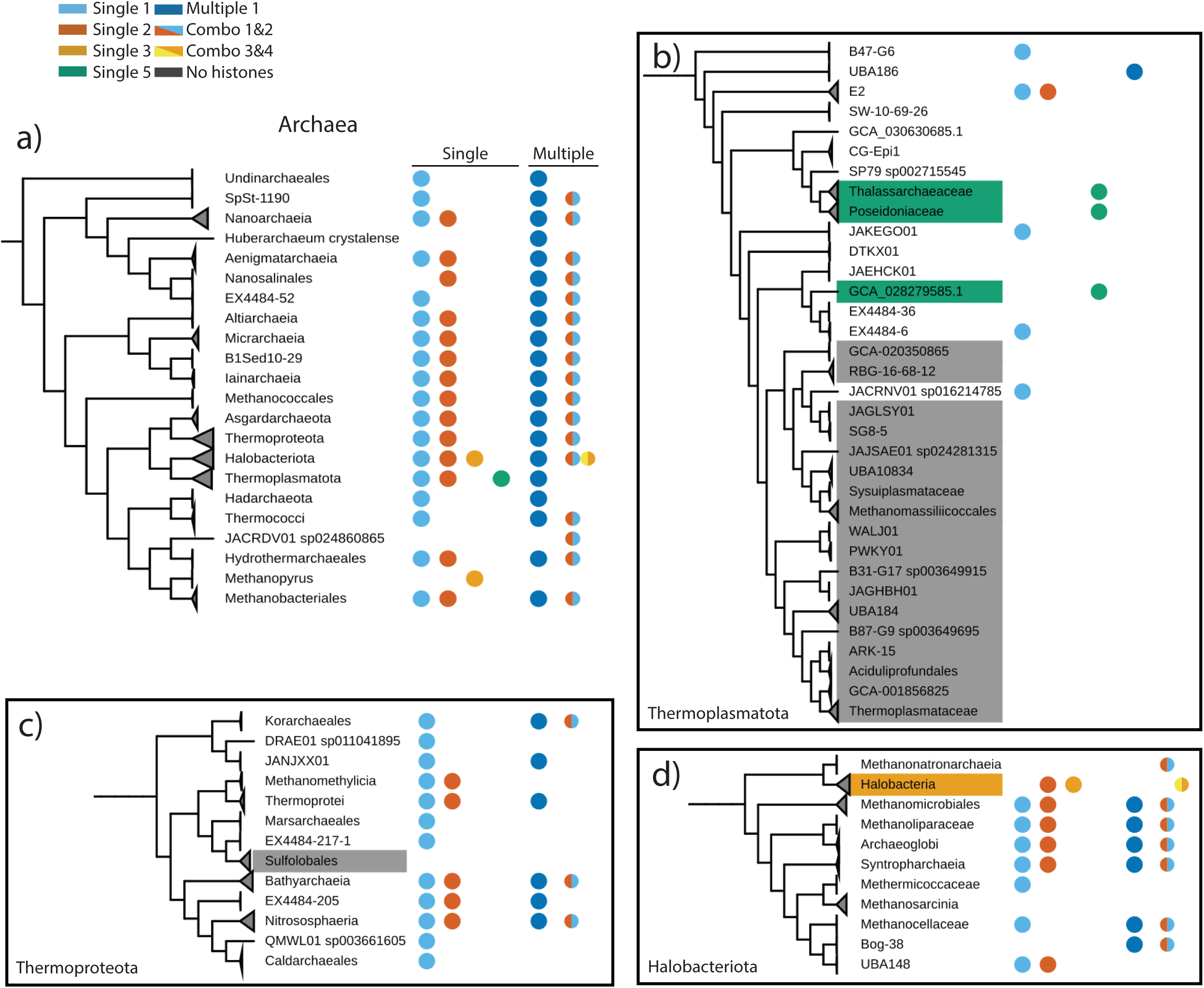
Some, but not all, strategies are taxonomically restricted. Colored dots indicate the presence of at least one genome in a taxon that employs a particular histone strategy. Histone clusters are colored as in Figure 1. **a**) Phylogeny of major archaeal groups. **b)** Phylogeny of Thermoplasmatota, where most taxa do not encode histones (highlighted in gray). Genomes encoding histones from cluster 5 are exclusive to the order Poseidoniales (families *Thalassarchaeaceae* and *Poseidoniaceae*) and a single closely related genome (all highlighted in green). **c)** Phylogeny of the Phylum Thermoproteota showing the absence of histones from the order Sulfolobales (highlighted in gray). **d)** Phylogeny of Halobacteriota showing the exclusive presence of cluster 3 with cluster 4 (acidic doublet with acidic miniature histone) in the class Halobacteria (highlighted in orange), and the near exclusivity of the combination of single basic and acidic doublet histone, except for the genus Methanopyrus). More details are provided in Supplemental Figure 4. An interactive and expandable tree containing histone annotations of the seven major strategies is available through iTOL: https://itol.embl.de/tree/1281386427164781716417921

Of the two most frequent combinations of histones, combination 1&2 (basic and acidic singlet) occurs in large groups in Methanobacteria and Halobacteria, and in smaller groups elsewhere in the domain (Figure 3c, Supplemental Figure 3). Combination 3&4 (acidic doublet and acidic miniature) is restricted to Halobacteria (Figure 3d). We also confirmed previous findings that histones are exceedingly rare in the class Thermoplasmata (formerly Crenarchaeota) or in the order Sulfolobales (Figure 3b, Supplemental Figure 3)^23^. A full list of histones and their corresponding genomes can be found in the supplemental materials.

### Selective pressures may influence strategy type

To understand the selective pressures associated with a specific histone cluster or strategy, we scoured metadata linked with the GTDB genomes for correlations. Specifically, we focused on genome size, GC content, coding density, and sampling location. Only two of the histone strategies (single 3 and combination 3&4) are found in organisms with genomes that are significantly larger, and have a higher GC content than those that do not encode histones (Figure 4a, b). This is probably because increased GC content is a known adaptation to high saline environments, and it is mostly halophiles that employ this strategy^35^. Protein coding density is somewhat higher in genomes encoding cluster 5 histones (single 5; Figure 4c). Despite these subtle differences, our analysis does not explain why a sub-group of archaea does not appear to employ histones.

**Figure 4:**
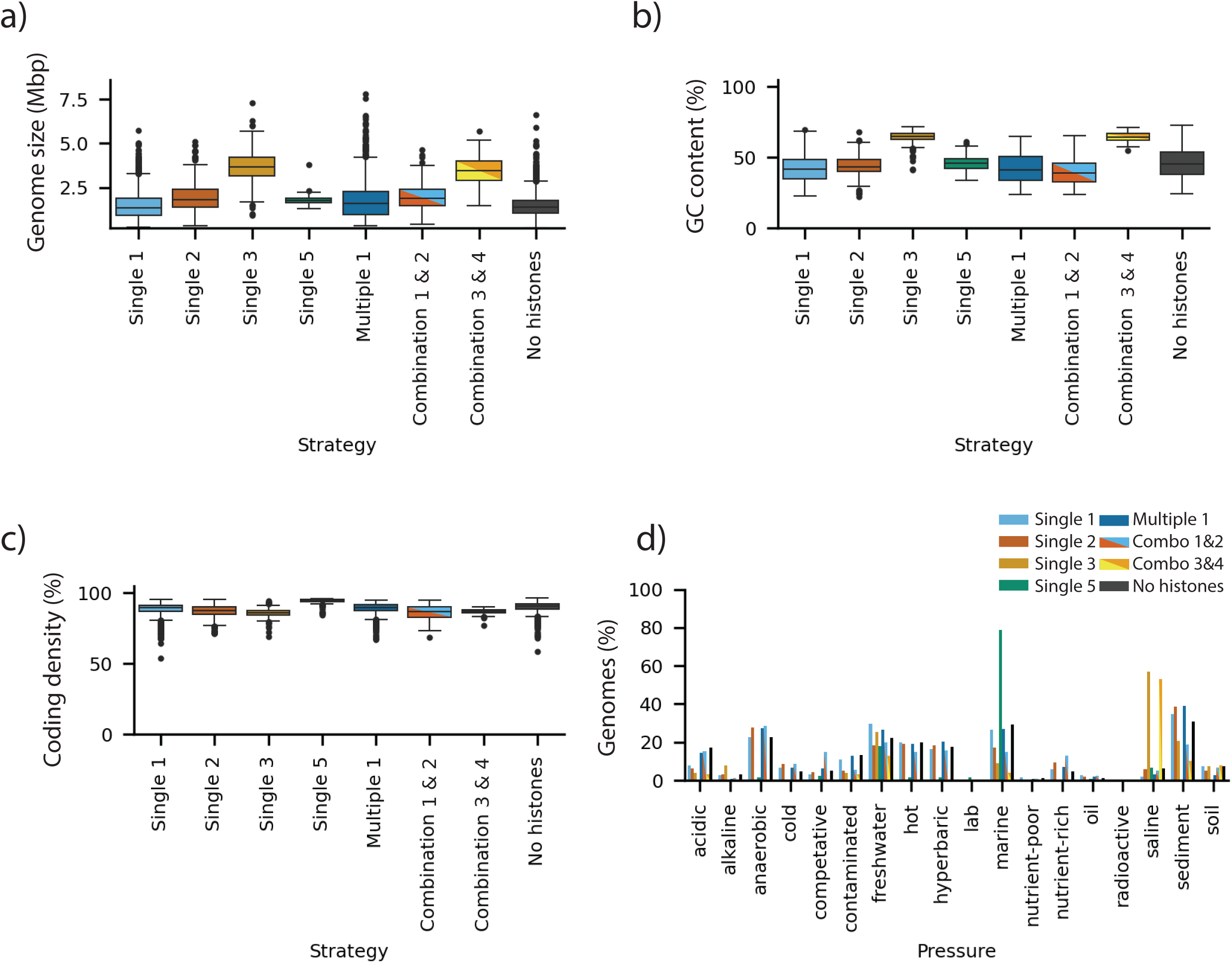
Correlation of histone presence and strategy with genome size, GC content, coding density, and environmental pressure. **a**) genome sizes. **b**) GC content. **c**) gene coding density. **d)** Histone strategy versus environmental pressures (Supplemental Table 2).

We also coded keywords found in genomic sample location annotations and found that some environmental pressures appear to correlate with specific combinations of histones (Supplemental Table 2). For example, archaea growing in anaerobic conditions tend to have combination 1&2 histones, and archaea found in extremely saline conditions seem to be enriched for combination 3&4 histones (Figure 4d, Supplemental Figure 4). It should be noted that these parameters are harder to quantify and verify, and as such the correlations have to be taken with caution.

### Sequence bias and conservation of archaeal histones

To better understand the differences in physical parameters between all archaeal histones, we compiled the overall composition of amino acids in each histone cluster (Supplemental Figure 5). There is an abundance of amino acids with a high propensity to form α-helices such as alanine, isoleucine, leucine, and valine, as expected for histones which are primarily α-helical. In the different histone clusters, we saw enrichment of, or bias away from specific residues compared to the overall sequence composition of all archaea. Notably, archaeal histones outside of cluster 1 have acidic isoelectric points (Figure 1c, Table 1). This is surprising as eukaryotic histones invariably have a positive overall charge and require basic residues (arginine and lysine) to effectively bind DNA in eukaryotes^36^. Besides an enrichment in acidic residues, the acidic histones from clusters 3, 4, and 5 exhibit the classic halophilic protein adaptation of a compositional bias from lysine to arginine^35,37^. Histones from these groups are also characterized by a higher percentage of the aromatic amino acids phenylalanine and tyrosine, which are both known to stabilize proteins in harsh environments (Supplemental Figure 5d,e,h)^37^. Across all archaeal histones, tryptophan and cysteine are underrepresented compared to the universal proteome, perhaps due to their metabolically expensive nature^38^.

Overall, archaeal histones are much divergent in their amino acid sequence than eukaryotic histones, which are among the most conserved proteins known^39^. The degree of conservation is particularly high in histones from halophilic organisms (cluster 3 and 4 histones) which could be due to a sampling bias towards closely related halophilic organisms, or due to a biological restriction to residues that facilitate function in hypersaline environments (acidic/aromatic residues, see above).

The cluster 1 and 2 histones that exist as the sole histone in the genome exhibit no characteristic sequence features compared to the histones that co-exist with others (Supplemental Figure 3, compare panel a. b with e, f, and g). Similarly, cluster 3 histones have the same sequence features whether they occur alone or with cluster 4 histones. Cluster 4 is interesting as the length of the ∼150 members is almost universally conserved to 55 amino acids and shares many of the characteristic features with the bacterial histone Bd0055^5^ (Figure 1b). Cluster 5 histones are unique in that they always occur as the sole histone-encoding gene, and their sequences are not well conserved. Aside from the first histone fold motif in these sequences, they do not contain many of the classic histone signatures (see below), suggesting that they may have co-opted the histone fold to perform a different function in the cell.

Specific ‘histone signature motifs’ are common to the majority of histones (boxed sequences in Supplemental Figure 6, and shown for HMfB in panel j). The ‘RKTV motif’ is located in the L2 loop connecting helices α2 to α3. While the first three amino acids in this motif (RKT) are present in nearly all archaeal histones, V is often substituted by I or L, but is universally a hydrophobic amino acid. In all known histone structures from all domains of life, this loop pairs with the less conserved L1 loop of the second histone in the histone fold dimer to form the L1L2 DNA binding motif^2^. In the L1 loop, the ‘RV motif’ is found throughout the majority of archaeal histones. Valine packs against the conserved hydrophobic side chain in the L2 RKTV motif to stabilize the underside of the paired L1-L2 loop, and the arginine extends into the minor groove of DNA and that is stabilized by a threonine in the RKTV motif (the RT pair). We also note the strong conservation of a salt bridge that stabilizes the L2 loop in its critical conformation (the ‘R-D clamp’), which involves the arginine in the RKTV motif and a conserved aspartate, invariably located 7 amino acids downstream in the α3 helix of the histone fold, even in the rudimentary α3 helix in cluster 4 histones (Supplemental Figure 6i). In eukaryotic, archaeal, and viral nucleosomes for which the structures are known, the fixed L1L2 configuration poises the main chain of both loops to contact the DNA phosphodiester backbone, and orients the arginine in the L1 loop (RV motif) to point into the compressed minor groove of the DNA. **As such, these conserved amino acids in the L1-L2 loops represent a universal histone signature in addition to the ability to form histone fold dimers that might be useful to identify other histone-like proteins.**

Unique to archaeal histones, a glycine in the L1 loop that we previously showed to be essential for hypernucleosome formation in *T. kodakarensis* histone HTkA ^4^ is also highly conserved throughout histone clusters 1, 2, 3 (for class 3, only in the N-terminal histone domain), but not in clusters 4 and 5. This suggests that histone clusters 1-3 might be able to form closely stacked hypernucleosomes.

### Structural prediction of histone complexes: histone homo- and heterodimers

To predict whether multiples of histones might be used to bind DNA and fold into nucleosome-like structures, we used AlphaFold3 to build models of a representative histone from each strategy as dimers or as tetrameters^40^. To choose unbiased representative histone candidates for each of the seven strategies, we calculated the center of mass in the four dimensions used in Figure 1 for each histone in each strategy from Figure 2 and identified the sequence closest to that point. For genomes that encode two histones, we chose a genome that encodes the histone closest to the center of mass of one of the clusters and used both histones from that genome as representatives (Table 2).

**Table 2.**
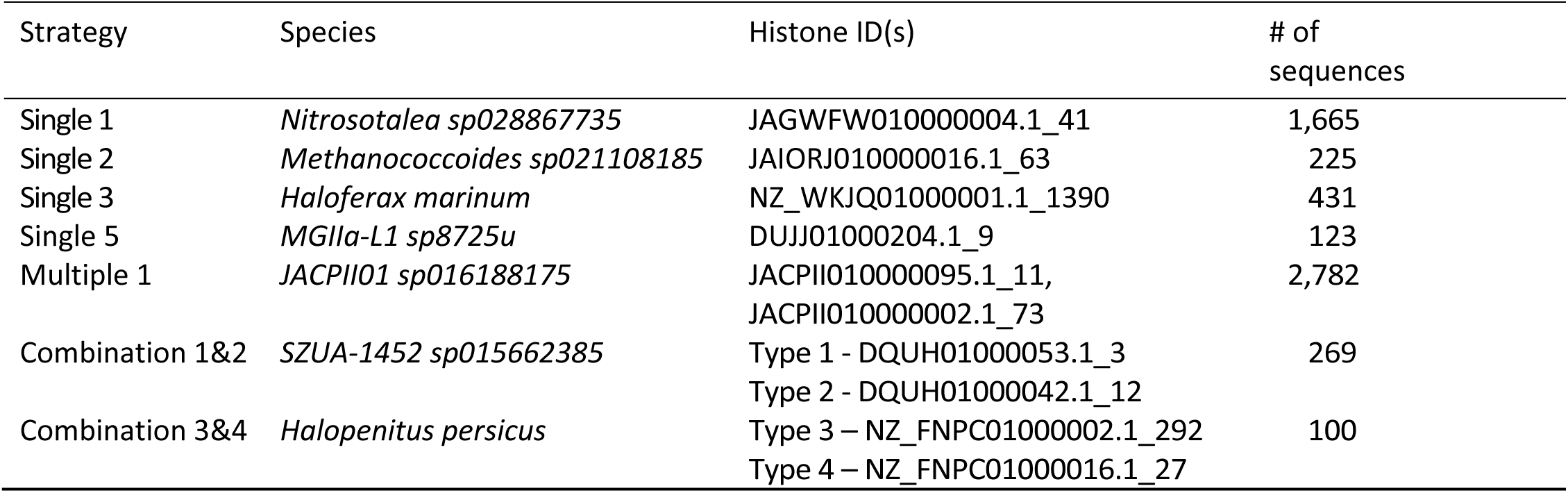
Representative genome for each strategy. Representative genomes were identified by encoding the closest histone to the average of all histones in each strategy, using the four physical parameters from Figure 1. For strategies with multiple histones, genomes were chosen by calculating the closest to the average histones for each type within the strategy, and then choosing the genome that had the most prevalent composition of histones for that strategy (two histones for multiple 1, one of each for combinations 1&2 and 3&4). All amino acid sequences are found in supplemental data.

The representatives from cluster 1, 2, and 3 form homodimers that resemble known structures of histone fold homodimers (Supplemental Figure 7). The N and C termini of single 5 histones can adopt conformations similar to histone fold dimers through intra- and interchain interactions, respectively.

Basic histones that occur in genomes together with other histones are likely able to form both homo- or heterodimers (as shown experimentally for *M. fervidus,* an organism that employs the multiple 1 strategy)^41^. In the median organism representing this strategy, the second histone is missing a well-defined α3 helix, yet this histone is also able to homo- and heterodimerize *in silico* (Supplemental Figure 7b). A number of histones from this cluster appear to have a prematurely terminated α3 helix, yet are still able to form homo- and heterodimers (not shown). In the median organism combining a basic and acidic singlet within its genome (combination 1&2), homo- and heterodimers are predicted with similarly high levels of confidence (Supplemental Figure 7c). It is important to note that even the models of clusters with acidic charge maintain a basic putative DNA binding ridge on their outer surface. In our representative employing the combination 3&4 strategy, combining one acidic doublet with one acidic miniature, the doublet folds into a structure that is very similar to a canonical histone fold dimer. The acidic miniature histone from cluster 4 is predicted to fold into a homodimer that has closer resemblance to the bacterial histone Bd0055 than to HMfA or HMfB.

### Some, but not all archaeal histone-fold dimers form tetramers via a four-helix bundle structure

The ability to form tetramers from histone fold dimers through well-defined four-helix bundle (4HB) structures is a hallmark of all canonical nucleosomes^3,4,6^. This interface is formed by the pairing of the C-terminal end of the long α2 helix and the α3 helix of two separate histone dimers (Supplemental Figure 8a, circled). We used AlphaFold3 to predict whether the histone fold dimers shown in Supplemental Figure 7 are capable of forming tetramers through the 4HB or any other means. Note that we display the solution with the highest level of confidence, with the acknowledgement that in some instances alternative solutions are created with only slightly less favorable IPTM scores (see, e.g. Supplemental Figure 8d, inset).

Representative histones from strategies using a single cluster 2 or 3 histone are all predicted to form homo-tetramers through canonical 4HB assemblies that resemble archaeal HMfB and eukaryotic histones H2B-H4 and H3-H3’^3,4^. Basic singlets from ‘single 1’ may form closed tetramers (as is the case for our median histone sequence) as well as open, canonical tetramers. Note that such closed tetramers have not been observed experimentally for any archaeal histone with the typical 28 amino acid long α helix, while open tetramers have been visualized in various complexes with DNA^4,30,31^. In our experience, these predictions have to be taken with a healthy dose of skepticism: for example, for HMfB, for which structures are known, AlphaFold3 predicts a closed and open tetramer as well as a ‘back-to-back tetramer’ that doesn’t involve the 4HB with closely spaced confidences, but generates an open tetramer resembling the experimentally determined structure when calculated in the presence of DNA. No combination of histone fold dimers from the single 5 representative is predicted to form higher order assemblies mediated by a 4HB (Supplemental Figure 8b, green).

Our representative basic histone that co-occurs with a second basic histone (multiple 1) is only predicted to fold tetramers from a homodimer of histone A. Histone B alone, or combined with histone A, does not form a tetramer *in silico,* but we did not explore whether this is a general phenomenon of histones from this cluster (Supplemental Figure 8c). Histones from combination 1&2 (basic and an acidic singlet; a wide-spread combination) are predicted to form open tetramers either from the basic histone alone, or from basic-acidic histone fold dimers (Supplemental Figure 8d). The acidic histone fold homodimer can form a tetramer through a variety of arrangements with nearly the same confidence (inset). Finally, the combination of an acidic doublet and an acidic miniature (combination 3&4), specific to and prevalent in halophiles, is not predicted to form a heterotetramer. While the acidic doublet forms an open ‘tetramer’ structure, the acidic miniature assembles into either ‘back-to-back’ (shown) or face-to-face tetramers with similar confidence.

### AlphaFold3 predictions and simulations suggests that most histones form stable structures with DNA

To explore whether these systems might form plausible complexes with DNA, we employed AlphaFold3 to predict structures with DNA and then evaluated their physical stability using all-atom molecular dynamics simulations. We predicted nucleosome models with the equivalent of 8 histone folds from each histone strategy and 147 bp DNA, sufficient for forming a nucleosome-like arrangement. All combinations that are able to form canonical, open tetramers via the 4HB are predicted to wrap DNA around the outside of the histone torus (Supplemental Figure 9), as is the representative that is predicted to form closed tetramers in absence of DNA (Supplemental Figure 9c, of note also the case for HMfB). Cluster 4 and 5 histones are not predicted to wrap DNA. Note that the IPTM scores are rather low for all models except for those with combination 1&2 (Supplemental Figure 9d).

We ran molecular dynamics simulations of all structures that formed nucleosome-like structures for 100 ns in triplicate, to allow for relaxation and sampling of conformational flexibility. Our goal was to determine whether the AlphaFold3 models shown in Supplemental Figure 9 are energetically plausible. In these simulations, the human nucleosome and the archaeal structure from HMfB (for which there is a structure on shorter DNA, PDB 5T5K) remained tightly wound and experienced little conformational change, as judged by minimal movement of DNA during the simulation (Figure 5b). The median representative histone from an organism employing the same strategy (multiple 1) also formed stable nucleosome-like structures. Acidic single 2 histones form plausible structures, although they seem somewhat destabilized compared to the structures formed by the basic histones. Structures predicted with the single 3 histone cluster unraveled, eventually losing the protein-protein interactions crucial to maintaining a tightly wound nucleosome. As the original median histone formed an open structure, we also simulated a second nucleosome of this type which was predicted to form a closed structure (shown in Supplemental Figure 9), but both simulations resulted in similar ‘final’ structures with open conformations. Nevertheless, even these acidic cluster 2 and cluster 3 histones maintain their interaction with DNA throughout the simulation. Remember that single 2 and single 3 strategies are mostly found in halophilic organisms and, as such, simulations should likely be performed at much higher ionic strength. Indeed, similar simulations in 2 M KCl resulted in structures that remained closed (not shown).

**Figure 5:**
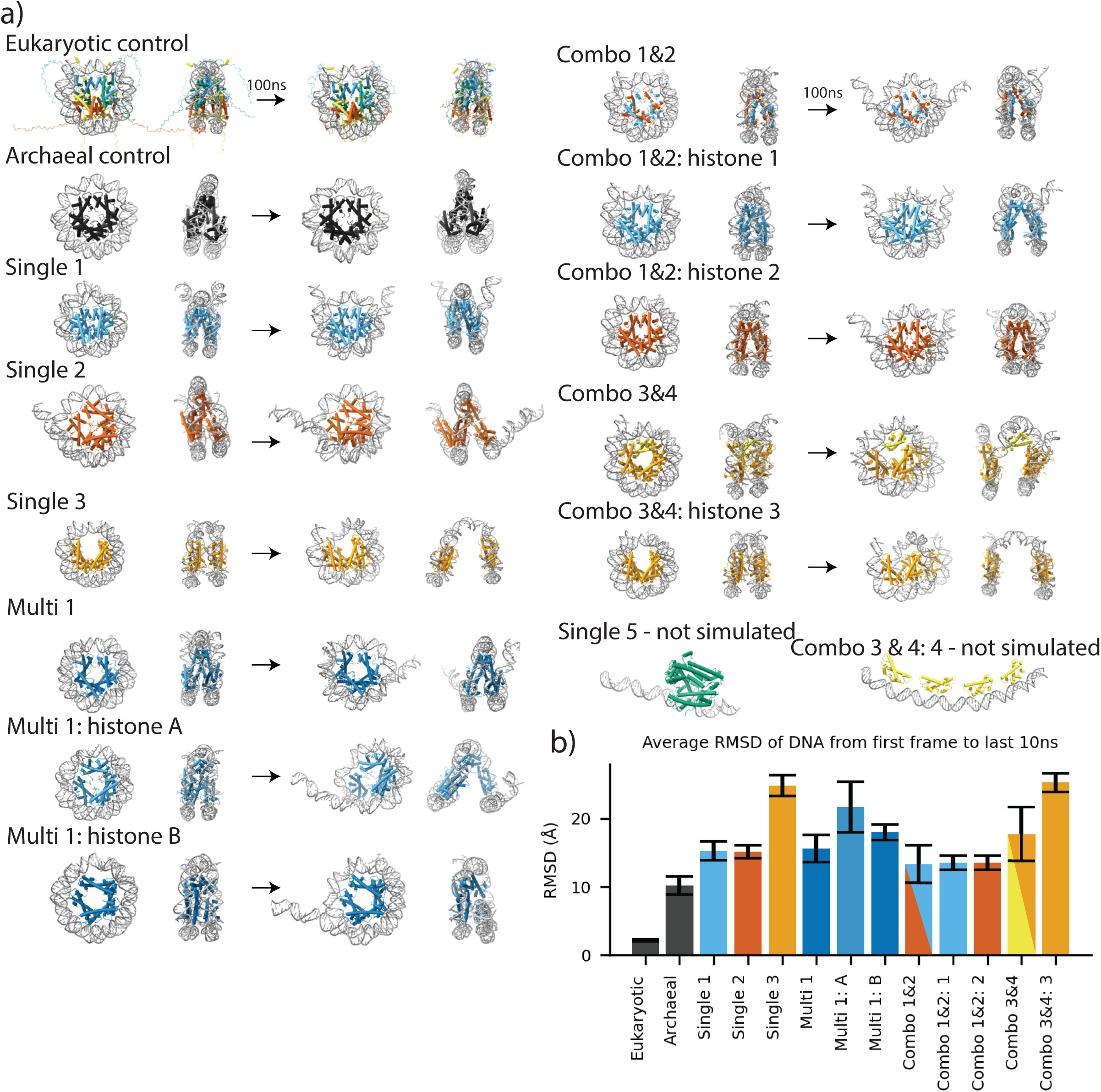
Molecular dynamics simulations suggest that not all of the predicted nucleosome-like structures are stable. All-atom molecular dynamics simulations of nucleosome-like particles predicted from each histone strategy (median representative, Table 2), in isolation or in combination. Simulations were started from an AlphaFold3 prediction shown in Supplemental Figure 9, using the equivalent of eight histone folds and 147 bp of Widom 601 double stranded DNA. AlphaFold models of the eukaryotic nucleosome and a nucleosome constructed from HMfA were used as controls. Simulations were run for 100 ns in triplicate. Side and face views of starting model and representative final model are shown. b) Change in DNA topology from beginning to end of simulation. RMSDs (Å) were calculated by averaging the RMSD of the DNA in each structure over the last 10 ns of the simulation to the starting frame. Because the DNA represents the topology of a nucleosome and is common to all the structures, we reasoned it was the most consistent way to monitor how much the models changed over time. Error bars represent standard error of the mean over three replicate simulations.

We also ran simulations of nucleosomes from genomes that encode multiple histones. We analyzed these histones in isolation and in combination (Figure 5), allowing us to query whether they might require a partner to form stable nucleosomes. For the two basic histones from the genome used to represent multiple 1 genomes, nucleosomes made from the combination of both histones or just histone B remained closed over the simulation. In contrast, nucleosomes made from just histone A appeared unstable (Figure 5a, b). This could suggest a means by which accessibility to the genome might be regulated, an idea that was recently supported by experimental data^31^.

As predictions were unable to assemble both histones from combination 3&4 into a nucleosome structure, we manually placed the cluster 4 histone (which co-occurs with cluster 3 histones) into the obvious space made when predicting the nucleosome with only cluster 3 histone dimers. We also simulated a nucleosome from cluster 3 histones in isolation. In the end, both simulation strategies resulted in unstable structures, except in one of the three combination 3&4 simulations where a wrapped conformation was maintained throughout the duration of the simulation. Whether this acidic doublet combines with the acidic miniature histone to form a nucleosome-like structure or not remains unknown. Likely, these structures require precisely oriented histones or high ionic stremgth to function properly, as organisms encoding them often utilize a “salt-in” strategy to cope with high levels of extracellular salt. The prediction of just the cluster 4 or cluster 5 histones alone did not result in nucleosome-like structures. If anything, these investigations highlight the limitations of AlphaFold3 in the prediction of histone assemblies beyond histone fold dimers, and emphasize the requirement for at least some degree of energy minimization.

## Discussion

Archaea are a diverse group of organisms that have adapted to a wide and extreme range of environments. With billions of years of evolution, this domain of life has diversified to meet unique challenges, and these adaptations presumably include strategies to protect and package genomes. This is particularly relevant for organisms that thrive at extreme temperatures, pH, and ionic strengths, all presenting challenges to genome integrity. While some archaea package their genomes exclusively with non-histone proteins such as Cren7, Alba, or Sul7d, the majority of them rely on histones^28^. Our search of available archaeal genomes reveals that 67% of known archaea encode histones that we group into five clusters depending on length, charge, hydrophobicity, and instability index. Because we used a rather conservative cutoff, it is likely that this percentage could be higher. The majority of histones are predicted to comprise a single, mostly tail-less histone fold domain with basic charge.

Four out of five clusters of archaeal histones (26.7 % of all histones) are acidic in character; they are either of canonical length (cluster 2), encode multiple histone folds in a single chain (cluster 3 and 5), or are predicted to have shorter α2 and α3 helices, resembling bacterial histones at least architecturally, if not in overall charge (cluster 4). Histones are encoded in genomes either by themselves or along with other histones, most commonly with one other histone of the same cluster, or combining two histones from different clusters. Intriguingly, 269 genomes encode an acidic and a basic histone that have < 60 % sequence identity, displaying a diversification in histone sequences that precedes the split into H2A, H2B, H3 and H4 that must have happened in early eukaryotes.

Using the structure prediction tool AlphaFold3, we predicted the structures of the ‘median histone’ for each cluster/strategy, in different oligomerization states and in complex with DNA. We deliberately chose the median histones rather than the nearest model organism to avoid bias and to best represent each cluster and strategy. Of note, many of these histones are derived from metagenomes and the corresponding organisms have not yet been cultivated.

Our analyses suggest that four out of the five clusters form canonical histone fold dimers, most of which tetramerize via a four-helix bundle (4HB) interface that is a hallmark of eukaryotic histone interactions. Representatives of clusters, either alone or in combinations, that tetramerize via a 4HB are predicted to organize DNA into nucleosome-like structures that remain stable in molecular dynamics simulations. These structures are similar to the experimentally determined structure of a cluster 1 histone in complex with DNA, which we showed forms a ‘hypernucleosome’ that may flex and open stochastically^4,30^. Histone-DNA interactions are maintained throughout the simulations for representatives of most clusters and strategies, even for those that have an overall acidic character. This is probably because even they maintain the ‘basic ridge’ around their outside that serves as a DNA binding surface. Of note, our simulations have not yet considered the diverse environments that our ‘median organisms’ might dwell in. For example, cluster 3 histones are mostly found in halophiles, and as such simulations at high (>2 M) ionic strength would be a better predictor of the plausibility of their histone-DNA complexes. Our data suggest that cluster 5 histones, even though they form plausible histone fold dimers, might not function in genome organization, and the role of cluster 4 histones (shorter acidic histones of highly restricted length to 55 amino acid, co-occurring with cluster 3 histones) remains unresolved. Importantly, given the limitations of AlphaFold (also demonstrated here), these predictions have to be verified experimentally.

While eukaryotes have largely selected for a narrow and conserved set of four histone sequences (plus a variety of histone variants^8^), archaea seem to be able to achieve genome organization with histones with much higher sequence diversity and using multiple combinatorial strategies. Nevertheless, the vast sequence space has brought into focus universal, functionally linked histone signatures, the RKTV motif and the R-D clamp in the L2 loop of the histone fold, the RV motif in the L1 loop, and the RT pair (Supplemental Figure 8j). In combination, these motifs serve to rigidify the L1L2 pairing to allow it to make main-chain interaction with the phosphodiester backbone of the DNA, and to orient an arginine to protrude into the compressed minor groove of DNA (referred to as a sprocket arginine)^42^. These signatures have been described over 25 years ago, and are reinforced here in a vastly expanded sequence space. The diversification in histone sequence outside of these motifs likely allows archaea to adapt to a diverse and extreme set of intracellular conditions than could not be tolerated by eukaryotic systems, and might afford them the ability to live in these environments without compartmentalizing their genomes.

Recently, other groups have used different tools to sample histone diversity across both archaea and bacteria. In a study by Dame and colleagues, histone sequences from archaea and bacteria were clustered into different groups based on sequence features^14^. This approach led the team to focus mainly on an array of bacterial histone sequences that are fused to other functional domains and whose functions are largely unknown. The work highlighted the power of approaches like HMMSearch to find disparate sequences which may fold into similar structures.

Our study emphasizes the need to explore these understudied and diversified classes of histones and to explore the biology of organisms that may otherwise be overlooked. By selecting organisms that broadly sample the diversity of archaeal histones, we can allocate resources strategically to maximize discovery. As many of the organisms have never been cultured, a logical next step to this work is to use structural biology and biochemistry to uncover how these histones physically structure DNA. An intriguing addition to the sparse availability of experimental structures has recently been published as a preprint, and suggests subtleties of archaeal chromatin structures that are caused by variations in histone sequence^31^. As recent breakthroughs in culturing (and, one would hope, genetically manipulating) archaea are revolutionizing the field ^43–46^, hypotheses gained from biophysical characterization could eventually be put to the test in the cell.

### Perspective

Our work highlights the power of structural prediction tools such as AlphaFold, yet demonstrate that they cannot (yet) replace experimental structures and biophysical analyses. To use these predictive tools properly, context and prior knowledge are necessary to avoid over-interpretation. For example, AlphaFold predicted the tetrameric structures of many histones to adopt conformations that appeared ‘closed’, yet when reinforced with a DNA sequence that is biased in the PDB to form nucleosomes, these same histones formed nucleosome-like structures. AlphaFold and similar tools are built on massive amounts of training data and usually do well when re-predicting structures they have trained on. Some models ignore the basic laws of physics, placing atoms on top of other atoms and predicting structures that fall apart in molecular dynamics simulations (Figure 5). At least for now, and for this system, the predictions are not yet ready to stand on their own without experimental validation, especially for the more complex models beyond histone tetramer, and in the presence of DNA.

## Methods

### Histone identification and HMMSearch optimization

Predicted archaeal protein sequences were downloaded from GTDB, release 220 (https://gtdb.ecogenomic.org/). This dataset included 11,277,496 proteins from 5,869 genomes, each with a specific taxonomic lineage. 7,140 putative histones were identified using an HMMsearch against PF00125 (PFAM for eukaryotic histones) and PF0808 (PFAM for archaeal histones). To establish which confidence thresholds to use with the JackHMMER and HMM-Search, we screened a range of expectancy values (E-Value) for each search strategy that went low enough to collect no hits and went high enough to be limited by filters built into the HMM algorithm (Supplemental Figure 1b). We noticed that most of the search strategies slowly collected hits up to an inflection point, where the number of hits began to increase rapidly. We reasoned that after this point the search models return mostly noise sequences. By iteratively clustering around this inflection point we were able to determine that hits above these E-values mostly constituted noise. Although most hits at the inflection point overlapped between search strategies, small outlier groups existed, so we combined the hits from both PFAMs around the inflection point and performed the rest of our analysis on this set (Supplemental Figure 1c). We eventually used E=4.0 for PF00125 and E=0.1 for PF00808.

### DBSCAN clustering

Histone sequences were imported with associated metadata from the GTDB. Ambiguous sequences were filtered out. Physical parameters of sequences were calculated using ProtParam from the Bio.SeqUtils python package. Histones were clustered with DBSCAN (implemented through the SciKitLearn package) optimized for a silhouette score of 0.25 (e=0.5 n=40) on the four parameters with the highest variance: length, pI, GRAVY, and helical propensity. Parameters were standardized prior to clustering using z-score normalization. To determine these clustering parameters, the data were randomly sub-sampled and tested with a range of parameters to optimize the silhouette score (Supplemental Figure 1). 0.25 was chosen as a target silhouette score, as it was able to reproduce clusters reliably after many rounds of clustering. After optimizing, the parameters were applied to two additional data subsets, verifying that the same number of clusters of roughly the same size were found in each. The physical parameters of each cluster were then calculated from the three subsets to define boundaries for the whole dataset. These ranges were tested on another three random subsets that were independently clustered to verify that the labels matched 95% of points in each test. The verified ranges were then used to label all points in the overall dataset. Proteins that failed to cluster into one of the five groups were removed from further analysis. Centers of mass and nearest neighbors were calculated for each cluster in standardized space and mapped back to real space. Edges were mapped linking histones coming from the same organism. Histones were then sorted into genomes and common strategies were calculated. Taxonomic data from GTDB was then used to map histone strategies onto a taxonomic tree using iTOL^47^.

### Metadata correlation

After assigning histones to strategies, a practical cutoff of 100 histones per group was applied to simplify analysis. Histones from groups which did not meet the cutoff were not used for further analysis, but were still included in the database. Metadata associated with each strategy were aggregated and comparisons to genomes without histones (No histones group) were preformed using the Shiparo test from the SciPy.stats Python package. We chose this test to deal with comparisons between datasets containing uneven variances. We chose metadata that we felt were most relevant to understanding the presences of histone: genome size, genomic GC percentage, and gene coding density.

### Environmental pressure correlation

We manually extracted the location data associated with each genome and coded keywords in each location to a set of standardized locations, which encompassed most of the genomes in the dataset. We then associated each of these locations with the environmental pressure(s) they most likely impart. A full list of keywords and coding can be found within the scripts.

### Sequence conservation

Conservation of histones from each group was calculated by taking the average occupancy at each position of aligned histones (aligned with MUSCLE) using a custom Python script^48^. ’Highly conserved’ residues represent residues whose conservation is at least one standard deviation greater than the mean conservation for that alignment. Conservations were calculated for each type of histone, both before and after strategies were assigned.

### Compositional bias

Amino acid composition of histone groups was calculated in Python using NumPy and plotted using Matplotlib. Composition was calculated on a per-residue biases, not as an aggregation of all the residues from all histones in a group. Because most histones in each group were of similar length, this normalization did not have a drastic effect, but still seemed appropriate to correct for a bias towards the composition longer sequences.

### Structural prediction

We predicted the structures of histones from each strategy as dimers (two histone folds), tetramers (four histone folds), and nucleosomes (with the addition of 147bp of dsDNA) using AlphaFold3, as implemented through the online server. We visually inspected each of the five models outputted by AlphaFold3 and proceeded with analysis on the highest confidence model not containing major clashes (usually the highest confidence model, i.e model 0). IPTM scores are reported in the figures.

### Molecular dynamics simulations

AlphaFold3 nucleosome predictions were used as starting models for simulations. Models were prepped for simulation using ChimeraX^49^. The terminal phosphate from each DNA strand was removed (to prevent simulation errors later), models were protonated, and then subjected to a few frames of MD implemented by using the ”Tug” function in ChimeraX and pulling on a single hydrogen atom at the terminus of a DNA strand. This ”Tug” step allowed the AlphaFold3 model to relax atoms and resolve clashes orders of magnitude faster than doing the same by hand. No gross topological changes were observed. All-atom molecular dynamics simulations with explicit solvent were carried out using AMBER and the ff14SB, bsc1, and tip3p forcefields (for protein, DNA, and water respectively)^50^. Structures were protonated again through TLEAP and hydrogen mass repartitioned in PARMED. Structures were placed in cubic boxes surrounding the structures by at least 25 °A, charge neutralized using potassium and chloride ions, potassium ions, and hydrated with water molecules. The structures were energy minimized in two, 5,000 step cycles: the first restraining the protein and DNA molecules to allow solvent relaxation and the second to allow full system relaxation. Minimized structures were then heated to 300 K and slowly brought to atmospheric pressure (1.01325 atm). The systems were then simulated for 100 ns in 4 fs steps. Simulation were performed in triplicate by starting the simulation over using a different random number during the heating phase. Distances between phosphates on neighboring residues at the center of a DNA strand were calculated as a proxy for nucleosome unfolding in representative simulations.

## Supporting information

all histone sequences

all histone sequences-parameters

interactive clustering plot

## Supplemental information

1. Supplemental tables 1 and 2
2. Supplementary figures 1-9
3. Interactive 3D chart displaying clustering data
4. Spreadsheet containing physical parameters of all archaeal histones
5. Spreadsheet containing all classified histones organized by genome

## Acknowledgments

his work utilized the Alpine high performance computing resource at the University of Colorado Boulder. Alpine is jointly funded by the University of Colorado Boulder, the University of Colorado Anschutz, Colorado State University, and the National Science Foundation (award 2201538)^51^. We also used the Blanca condo computing resource at the University of Colorado Boulder. Blanca is jointly funded by computing users and the University of Colorado Boulder^52^.

## Ethics approval and consent to participate

Not applicable

## Consent for publication

Not applicable

## Data availability

All underlying protein sequences and metadata were collected from <u>GTDB</u>. Histone sequences are provided in source data (excel spreadsheets).

## Materials availability

Not applicable

## Code availability

Code used to do analysis and run simulations is available on GitHub: https://github.com/shla9937/archaeal histone diversity

## Author contribution

SL conducted the analysis with input and editing from KL. SL and KL wrote the manuscript.

## Supplemental Material

**Supplemental Table 1.**
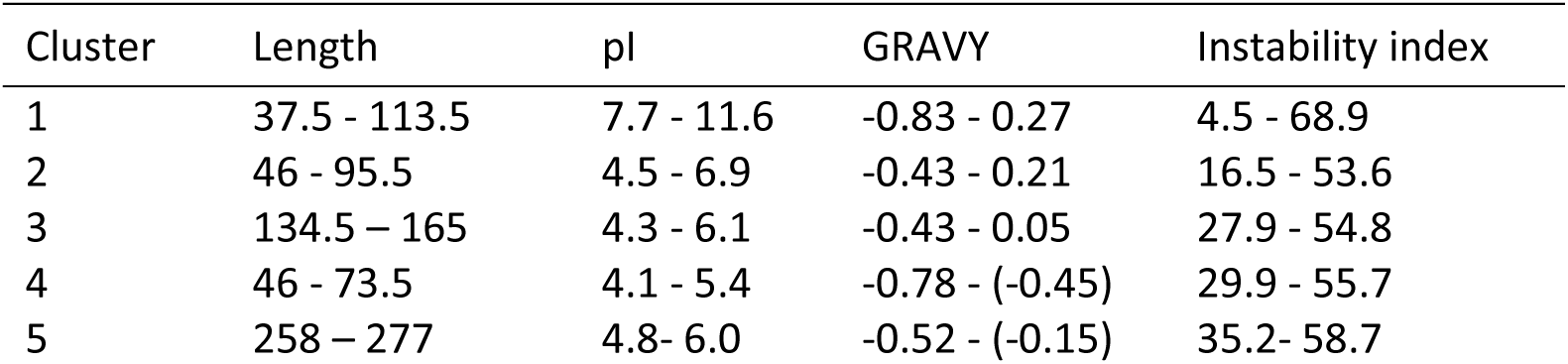
Ranges collected from DBSCAN clustering used to label histones. Ranges of physical parameters used to define each histones cluster in Figure 1. Physical parameters used are protein length (residues), isoelectric point (pI), hydrophobicity (GRAVY), and instability index.

**Supplemental Table 2.** Keywords used to parse genomic sampling locations from GTDB data.

| Keywords in location data | Location label | Associated pressure(s) |
| --- | --- | --- |
| acid, ph 4, ph 3 | acid | acidic |
| soda, alkaline | alkaline | alkaline |
| seep | cold seep | hyperbaric, cold, anaerobic |
| permafrost | permafrost | cold |
| salt, brine, saline | salt | saline |
| hot spring, hot pool, volcano, hot solfataric spring, geothermal | hot spring | hot |
| mine, mining | mine | contaminated, acidic |
| gut, intestine, rumen, ruminant, caecal, sheep, intestinal, caecum | gut | acidic, competitive |
| vent, hydrothermal, chimney, black smoker | hydrothermal vent | hyperbaric, hot, anaerobic |
| nuclear, olkiluoto | nuclear | radioactive, contaminated |
| sediment, mud | sediment | sediment |
| compost | compost | nutrient-rich |
| anaerobic digester, bioreactor, digestion, biodigester, reactor, fermentation | bioreactor/digester | nutrient-rich, anaerobic |
| waste, wwtp, sludge | waste | nutrient-rich, anaerobic, contaminated |
| landfill | landfill | nutrient-rich, anaerobic, contaminated |
| feces, fecal, stool, faeces, manure | feces | nutrient-rich, anaerobic |
| mouth, oral, dental | oral | competitive |
| estuary | estuary | marine, freshwater |
| ocean, marine, sea, atlantic, intertidal, tide-pools, bay, deep, depth, coastal, pacific | marine | marine |
| soil, grass | soil | soil |
| groundwater, well, aquifer, spring water | groundwater | freshwater |
| (not marine) + water, aquatic, wetland, river, pond, peatland, lake | water | freshwater |
| biofilm | biofilm | competitive |
| oil, petro (but not with soil) | oil | oil |
| lab | lab | lab |
| frac | fracking | oil |
| rock, aspo hrl | rock | nutrient-poor |
| none, metagenome, "" | none | none |
| (fallback case) | other | other |

**Supplemental Figure 1:**
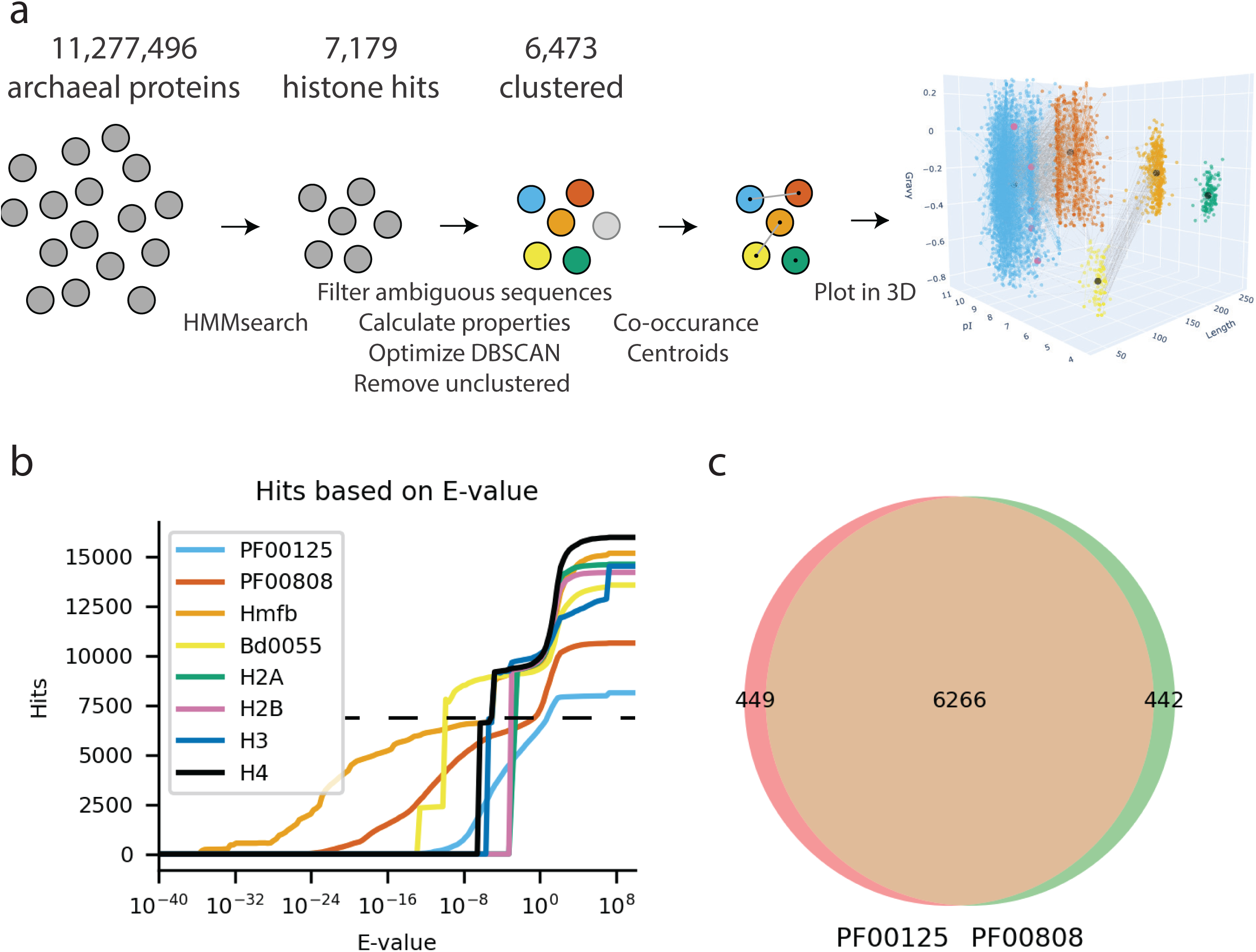
Iterative clustering strategy to generate Figure 1. **a)** All archaeal protein sequences from the GTDB (v220) were collected and selected for homology to archaeal and eukaryotic histones using HMMsearch against various inputs (outlined in b). Matching sequences were then filtered, physical protein parameters calculated, clustered using DBSCAN, and then unassigned sequences were removed. Co-occurrence networks (proteins in the same genome) were defined and centroids for each cluster were calculated based on nearest neighbor to center of mass of each cluster. The resulting data were plotted against three of the four features used to cluster them and colored according to cluster (1 – blue, 2 – red, 3 – orange, 4 – yellow, 5 – green). **b)** Number of hits obtained for a given E-value for HMMsearch (PF00125 and PF00808) or JackHMMer (Hmfb, Bd0055, H2A, H2B, H3 and H4 against all archaeal protein sequences. The inflection point denoted by the dashed line gave reproducible clusters when clustered over three randomly sampled subsets and was used to define histone hits for analysis. **c)** Overlap between histone sequences returned by HMMsearch using PF00125 (eukaryotic histones) and PF00808 (archaeal histones). Although there is a high degree of overlap, ∼440 were unique to either archaeal or eukaryotic histone searches. Combining these two datasets resulted in robust clustering and captured the majority of diversity observed in the other search strategies.

**Supplemental Figure 2:**
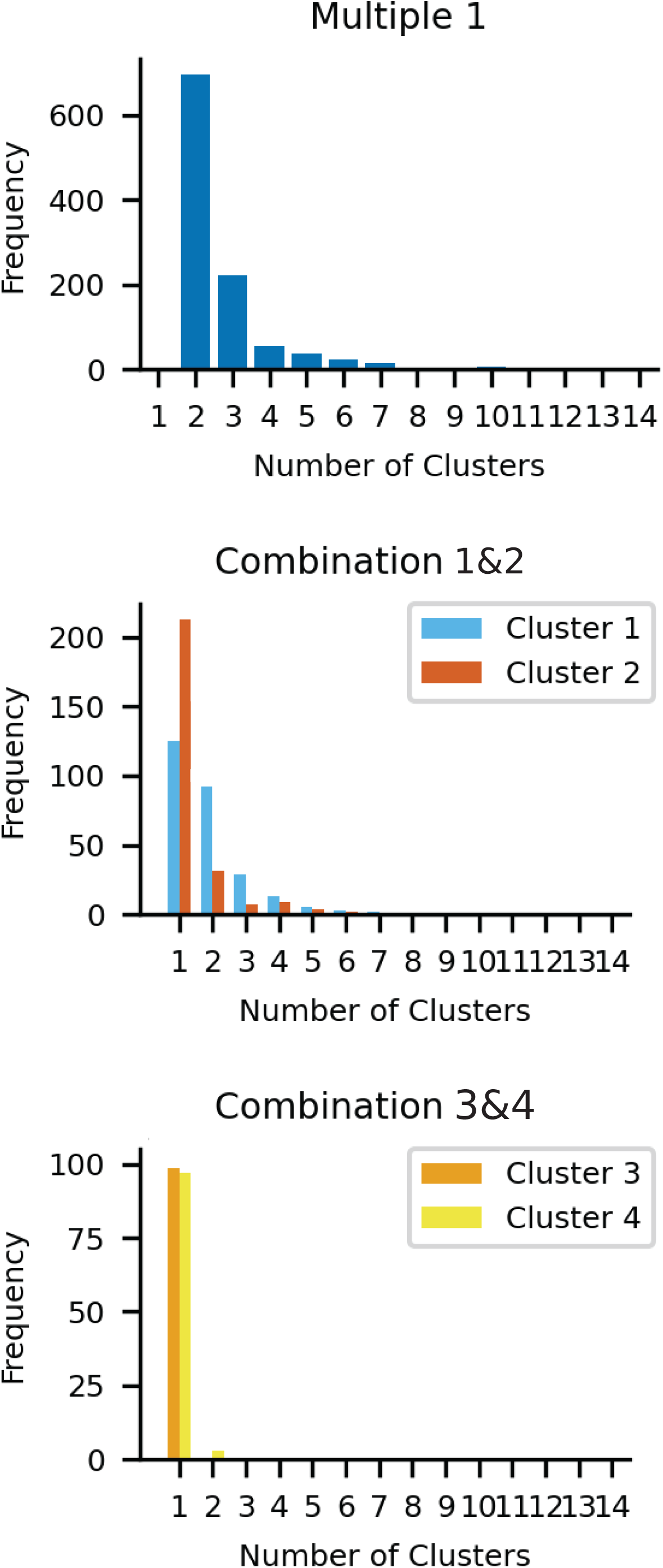
Breakdown of the number of each type of histone within genomes encoding multiple histones. Most genomes with multiple histones encode two. Combination 1&2 genomes often contain unequal ratios of cluster 1 to cluster 2 histones. Genomes from combination 3&4 encode cluster 3 histones at a 1:1 ratio with cluster 4 histones, except for three cases where only two cluster 4 histones are present.

**Supplemental Figure 3:**
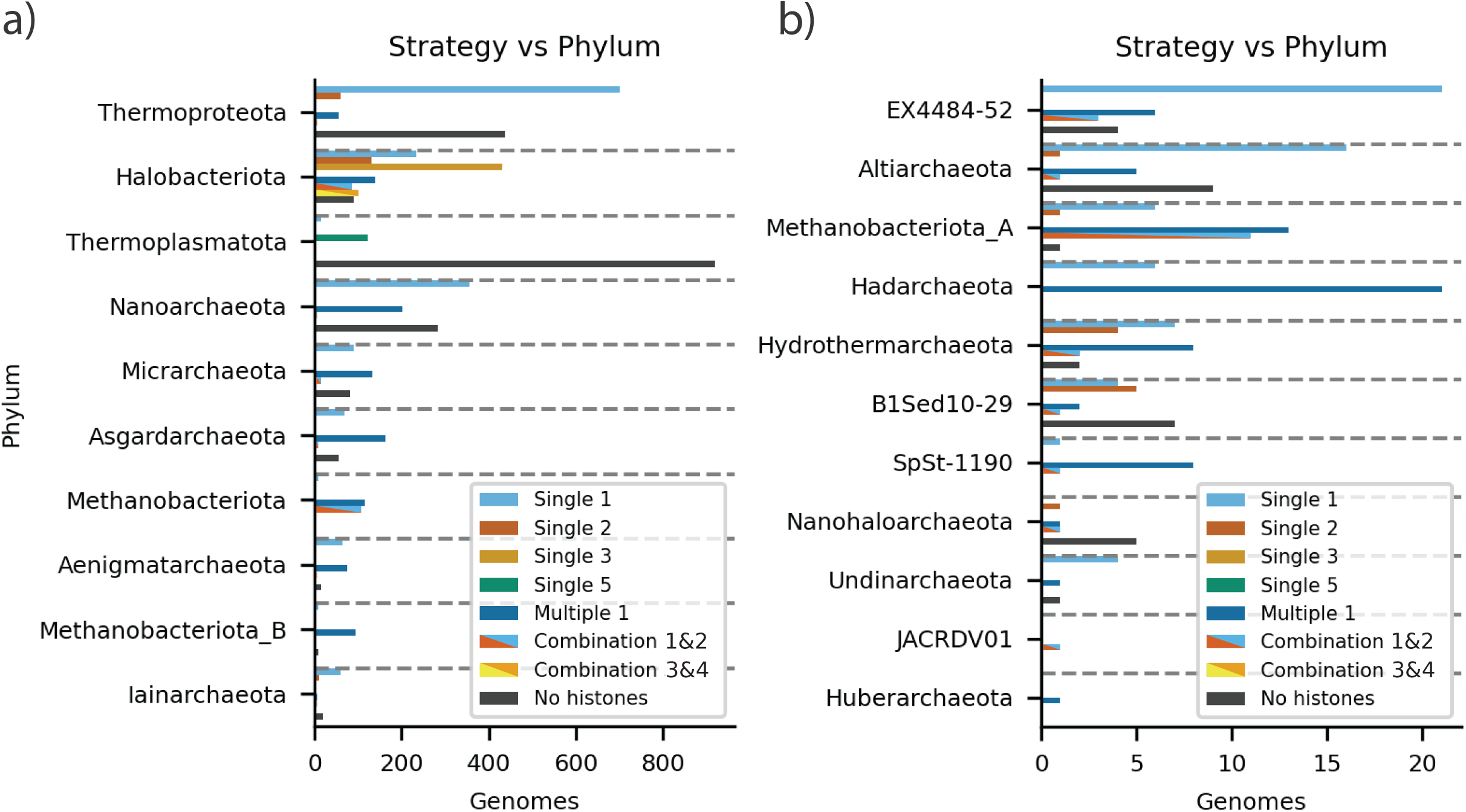
Number of genomes from archaeal phyla across employing a particular histone strategy. **a)** phyla which are represented by a large number of genomes. **b)** phyla with more sparse representation (note difference x-axes scale).

**Supplemental Figure 4:**
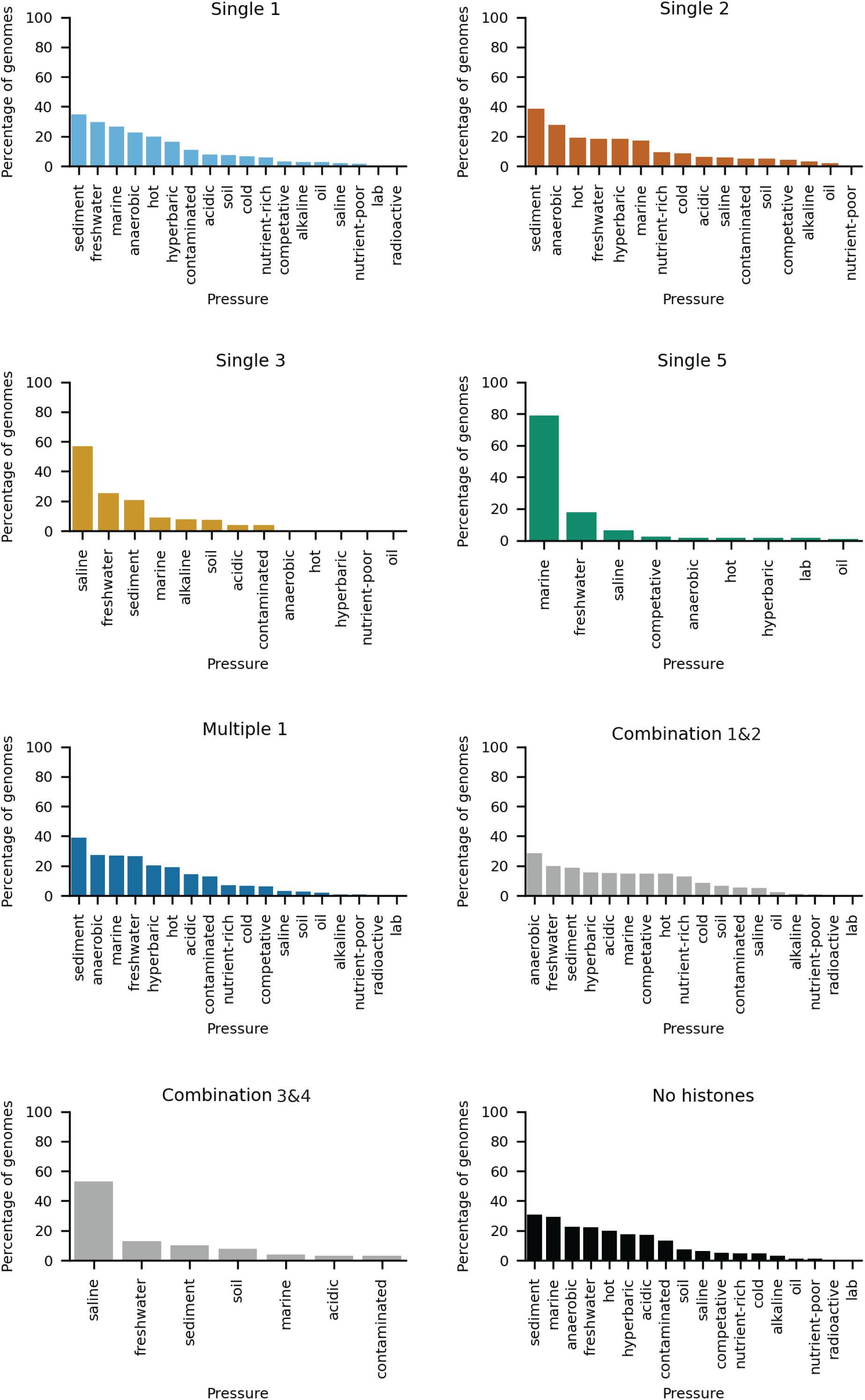
Histone strategy correlates with environmental pressure. Sampling locations for each genome in a strategy were curated from metadata and coded to common environmental pressures, then plotted as the percentage of genomes from that strategy that are associated with that pressure. Locations can be associated with multiple pressures. Pressures are ranked from most prevalent to least. Single 3 and Combination 3&4 showed a slight bias towards genomes from saline environments. Single 5 genomes are biased towards marine environments. Combination 1&2 genomes correlate with anaerobic environments.

**Supplemental Figure 5:**
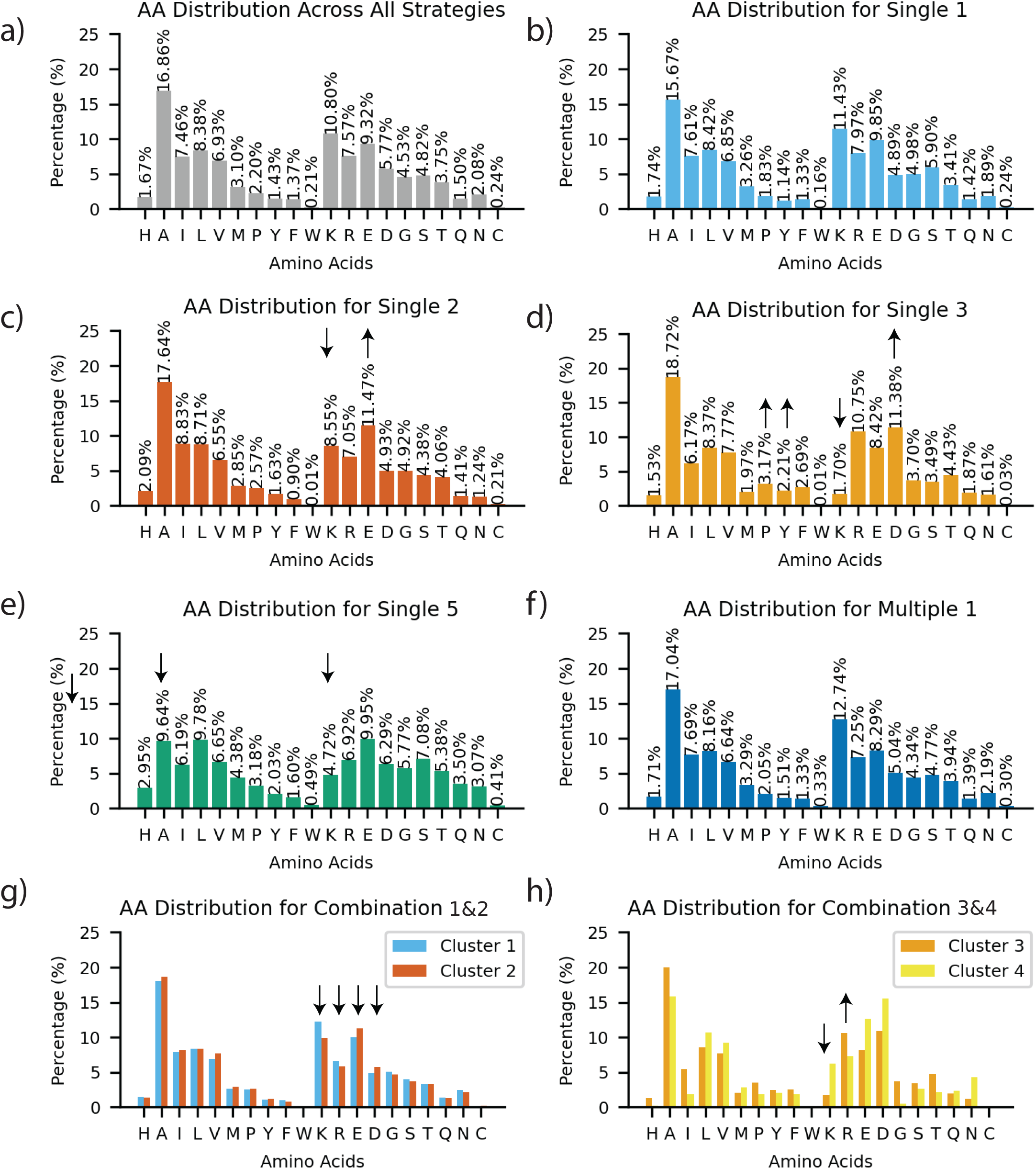
Amino acid composition of histones, grouped by strategy. Contribution of each histone is normalized for length, so that larger proteins do not dominate the average composition. Overall, archaeal histones are enriched in small, hydrophobic residues and basic residues. Tryptophan and cysteine are rarer compared to the ‘universal’ proteome, likely due to their metabolic burden. Arrows denote enrichment or depletion of a particular genome compared to the generic archaeal amino acid distribution. **a)** All archaeal histones; **b)** basic singlets (single 1); **c)** acidic singlet (single 2), arrows denote the shift in composition from lysine to glutamate, responsible for their acidic character. **d)** Acidic doublets (single 3). Arrows denote the increase of aromatic residues tyrosine and phenylalanine, as well as a marked shift away from lysine in favor of arginine and an enrichment in aspartate over glutamate. **e)** Single 5 histones, arrows denote the decrease in abundance of alanine and lysine. **f)** Multiple cluster 1 histones. **g)** Combination 1&2 histones. Arrows indicate the increase of basic and reduction of acidic residues in cluster 1 histones (blue) over cluster 2 histones (red). **h)** Combination 3&4 histones. Arrows indicate the increase of arginines and decrease of lysines in cluster 3 histones over cluster 4 histones.

**Supplemental Figure 6:**
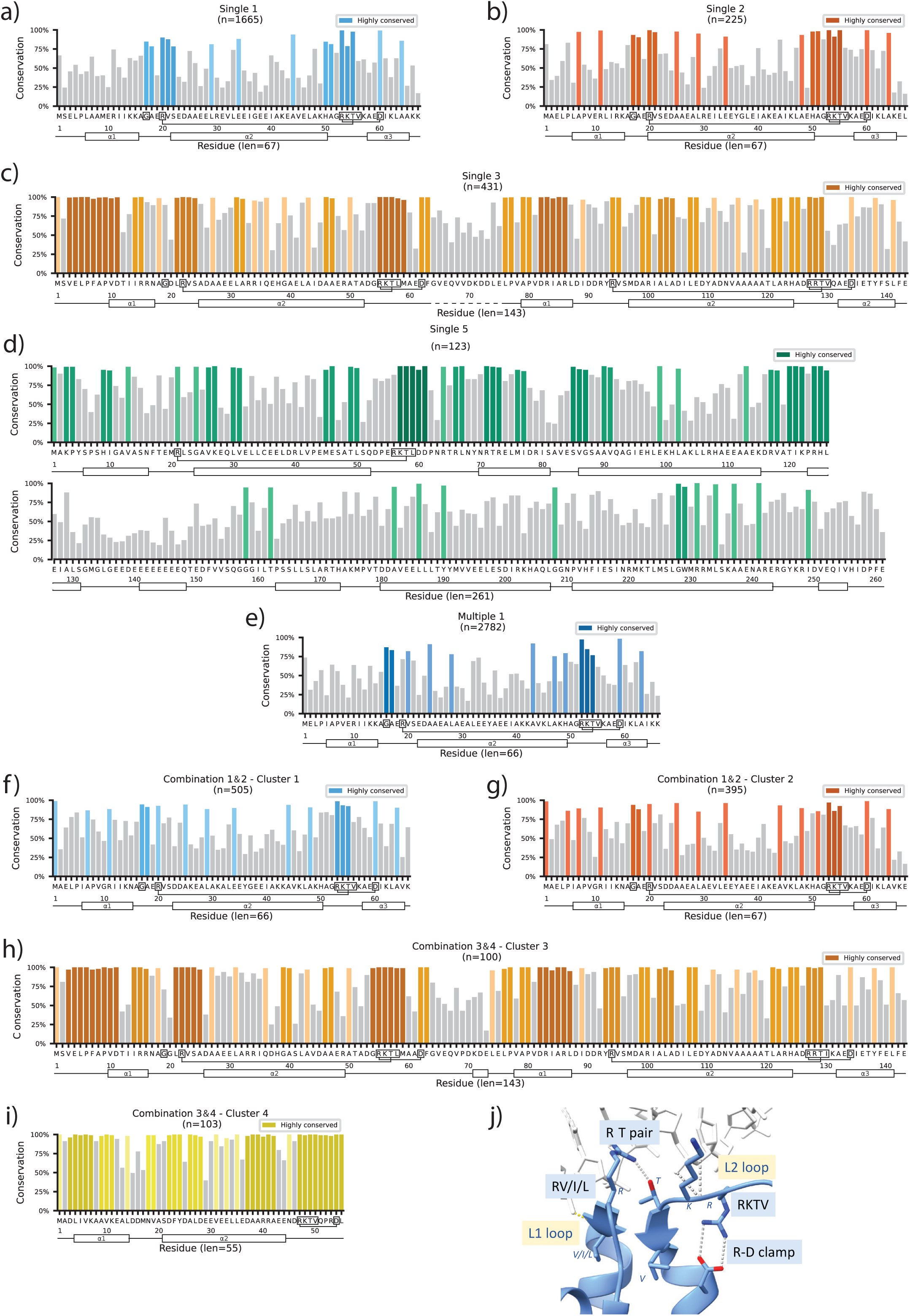
Conservation of histones by strategy. Amino acids that are conserved by more than one standard deviation than the mean conservation of an alignment are highlighted in color. Continuous clusters of residues having greater than one standard deviation of conversation are shaded in darker color for emphasis. Number of sequences in each alignment is denoted for each panel as “n=”. The average alignment length is denoted by “len=”. Conserved sequence motifs (RKTV motif, R-D clamp, RT pair, and G are boxed in the sequence, H-bonds for R-DNA clamp and RT pair are indicated). sPredicted secondary structure designation (from Supplemental Figure 7) are shown to indicate histone fold elements. **a-d)** singles, **e-i)** combinations, **j)** shows the L1L2 loop with conserved features, as indicated in the sequence alignments (pdb 5T5K).

**Supplemental Figure 7:**
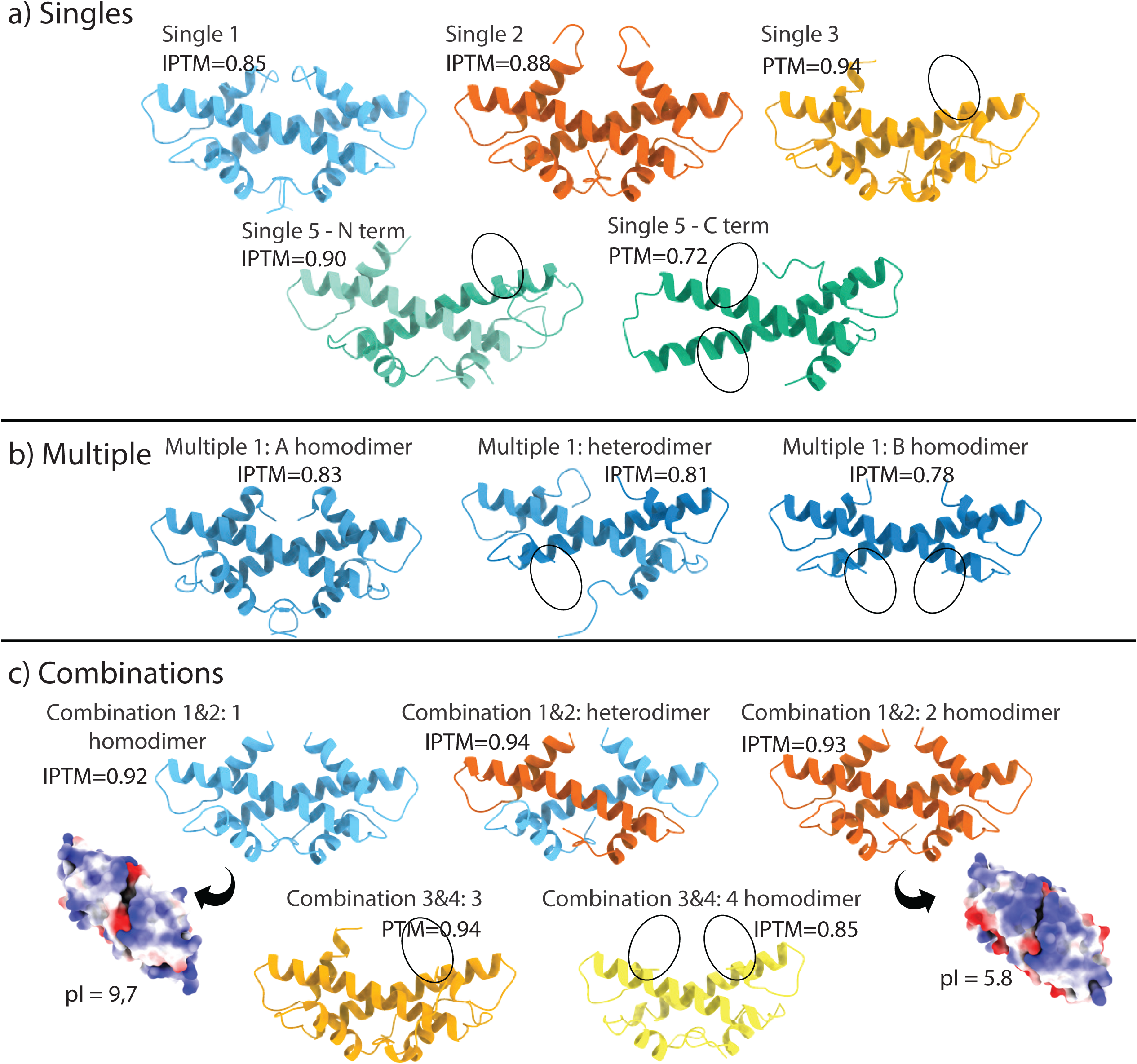
AlphaFold3 predictions of histone dimers. Models predicted from representative (‘median’) histone fold domains from each histone strategy (Table 2). Helices missing from the classic three-helix histone fold motif are indicated by circles. **a)** Histone dimers from representative organisms with only a single histone gene. For the single 5 histone, which contains five histone fold domains, the N-terminal two histone folds are split into separate chains and predicted as if belonging to two separate chains, whereas the C-terminal two folds were predicted as a single chain. **b)** Prediction of homo- and heterodimer histone fold structures from a representative of the multiple 1 strategy. **c)** Combination 1&2: basic and acidic histones can form homo- and heterodimers. Charged surface representation of the histone binding ridge are shown to the side of each homodimer Combination 3&4 histones were not folded together, as cluster 3 histones links two histone folds together in one polypeptide chain. All predictions have a high confidence score (IPTM).

**Supplemental Figure 8:**
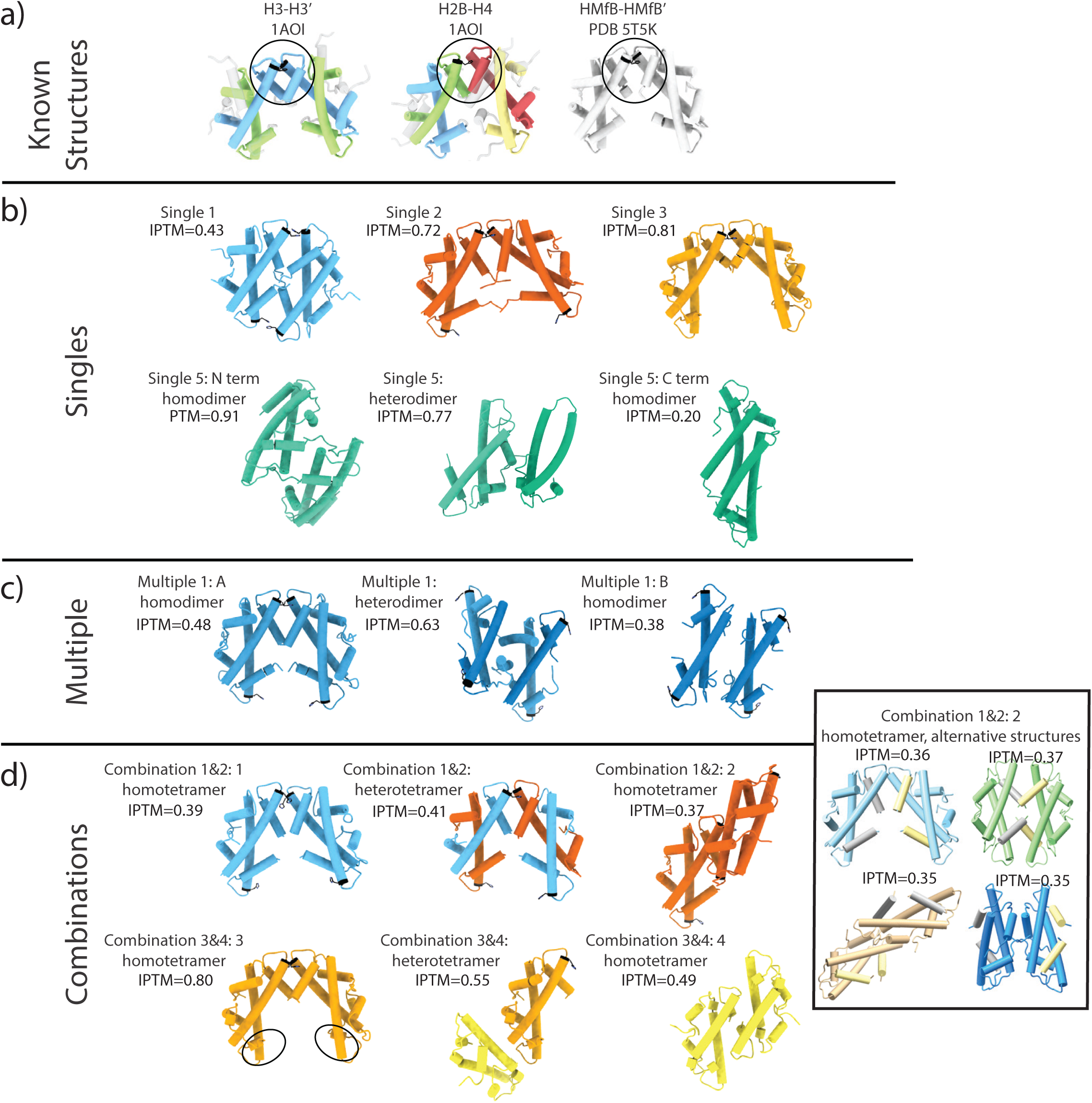
AlphaFold3 prediction of representative histone tetramers. Models predicted from four histone fold domains for the median sequence of each histone strategy. a) 4HB structures from experimentally determined structures. The conserved histidine is shown in black. **b)** Predicted homo-tetramers from single histone strategies. The Single 1 histone can form either closed (as shown here) or open tetramers with similar levels of confidence. Tetramers from the Cluster 5 histone do not appear to form canonical histone tetramers via four-helix bundle structures. **c)** Homo-tetramers from the multiple 1 genome vary in their predicted ability to form canonical open histone tetramers. **d)** The acidic histone from combination 1&2 forms canonical histone tetramers if paired with its basic partner. In isolation, it is predicted to fold into four completely different tetramers with similar confidence (inset; α3 helices are colored in grey and yellow for histone fold dimer 1 and 2, respectively; for orientation). The cluster 3 histone from combination 3&4 in isolation forms an open tetramer, but does not combine with its cluster 4 partner. The cluster 4 histone in isolation is predicted to form ‘back-to-back’ structures (shown), as well as closed tetramers with similar confidence.

**Supplemental Figure 9:**
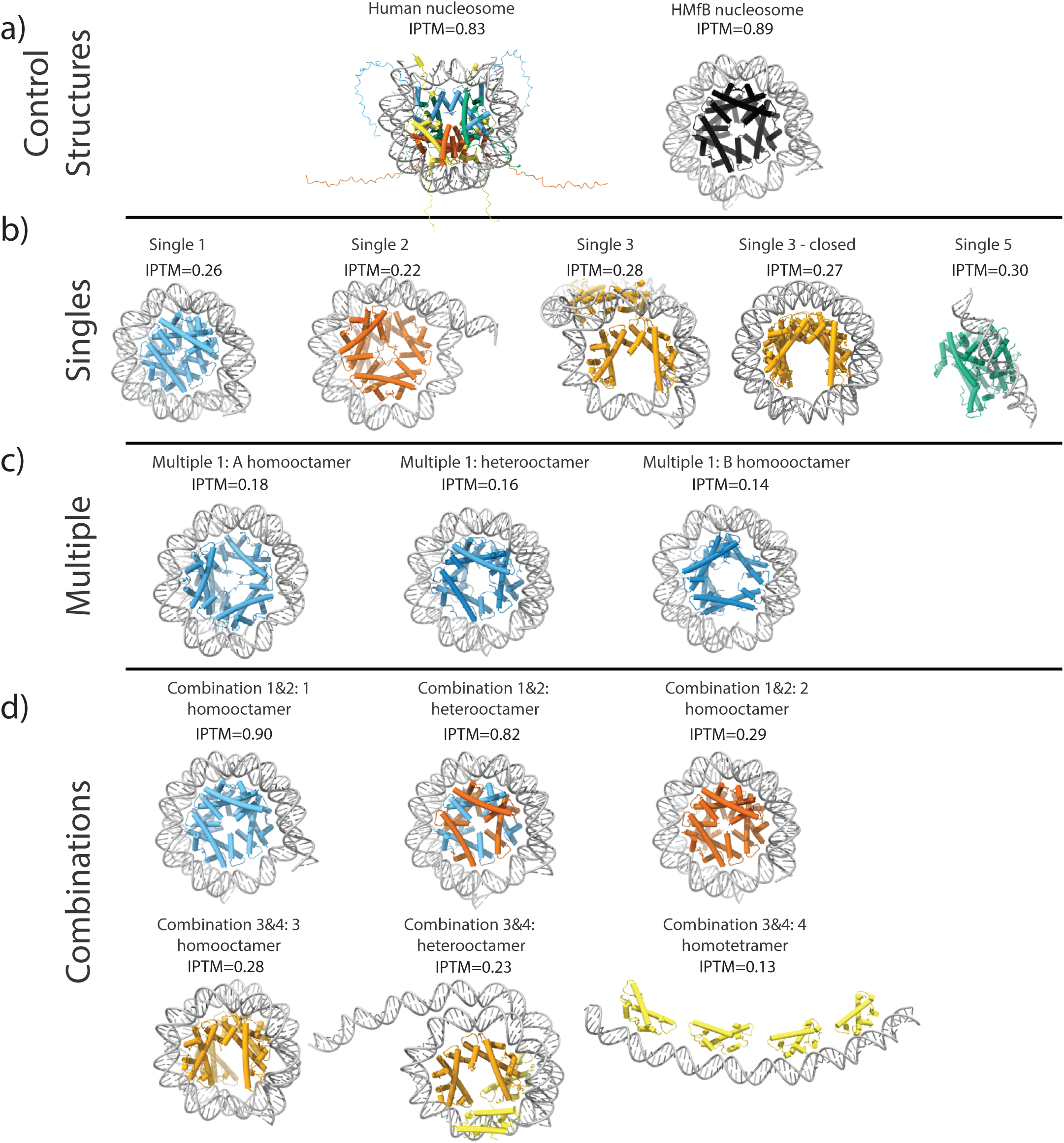
Predicting histone-DNA complexes with AlphaFold3. Models predicted from eight histone fold domains for a representative from each histone strategy, and 147bp of a nucleosome positioning sequence (‘601’ Widom DNA sequence). These serve as the starting structures for the simulations shown in Figure 5. **a)** Prediction of control structures of human and *M. fervidus* nucleosomes, closely resembling experimentally determined structures. **b)** Predictions for a basic and acidic singlet, and for the acidic doublet. For single 3, a second structure was predicted of a closely related histone (NZ_A0AIB010000141_204) that formed a closed nucleosome structure. **c)** the representative histones from the multiple 1 strategy appear to form nucleosome-like particles in each combination. **d)** Basic and acidic histones that co-exist in one genome fold into nucleosome-like structures either individually or in combination. In contrast, the acidic doublet does not combine with the acidic miniature, which on its own does not wrap DNA, nor does single 5.

